# Dual Metabolic Compensation to Dietary Choline Deficiency by the *Drosophila* Hologenome

**DOI:** 10.64898/2025.12.11.693343

**Authors:** Yuka Fujita, Satoshi Morozumi, Hina Kosakamoto, Makoto Arita, Fumiaki Obata

**Author notes:** These authors contributed equally to this work (Yuka Fujita, Satoshi Morozumi).

## Abstract

Coping with an unbalanced diet is critical in animals. However, the mechanisms that mitigate deficiencies of specific nutrients, such as choline, remain poorly understood. Choline is converted to phosphatidylcholine (PC), a major phospholipid. Using genetic, nutritional, and microbiome manipulation tools in *Drosophila melanogaster*, we identified two independent mechanisms that ameliorate the effects of choline deficiency. First, a strain of commensal bacteria synthesizes PC via choline-independent pathway and provides it to the host. Second, a host-derived phospholipid, β-methylphosphatidylcholine (βmPC), is generated from carnitine, which compensates choline deficiency as a functional analog of PC. Our findings reveal microbial and host metabolic processes that modulate dietary requirement for choline. This study also provides evidence that hologenomic coordination of metabolism enables animals to adapt to a micronutrient deficiency.

## Main Text

Animals have evolved diverse mechanisms to survive periods of nutrient scarcity. Numerous morphological, behavioral, and metabolic adaptations have been documented across the animal kingdom(*1–3*). However, many studies have focused on responses to general decrease in energy, or deficiencies of macronutrients, leaving the mechanisms that addresses the lack of individual micronutrients largely underexplored.

Choline is an essential nutrient whose dietary intake is required to sustain physiological levels, and its deficiency has detrimental effects on organismal health (*4*). Choline is a precursor of phosphatidylcholine (PC), one of the most abundant phospholipids, along with phosphatidylethanolamine (PE)(*5*). PC is indispensable for the integrity of cellular and lipoprotein membranes, and choline deficiency has been shown to inhibit growth and induce organ dysfunction, particularly in the liver (*4*, *6–9*). Accordingly, choline-deficient or methionine–choline-deficient diets are widely used experimental models for disrupted lipid metabolism and nonalcoholic fatty liver disease (*10*). Since phospholipids exhibit distinct physicochemical properties, organisms tightly maintain phospholipid homeostasis(*11*). Perturbation of this balance, such as through choline deficiency, can drive global shifts in lipid metabolism. For example, maintaining an appropriate PC:PE ratio through genetic manipulation mitigates the physiological consequences of choline deprivation in rodents (*12*, *13*). On a cellular level, modifying phospholipid acyl-chain length also alleviates the defects associated with low PC availability (*14*, *15*). Therefore, choline deficiency induces widespread remodeling of lipid metabolism, but whether this reflects coordinated adaptive responses or simply a direct consequence of reduced PC remains unclear, highlighting the need to dissect these mechanisms individually.

Given that choline deficiency alters lipid metabolism in ways likely influenced by the availability of other nutrients, isolating its specific effects demands strict control of overall dietary composition. To achieve this, we employed *Drosophila melanogaster* as a model organism. *Drosophila* can be reared on a chemically-defined synthetic diet, prepared from purified ingredients (*16*, *17*). This approach allows the dissection of organismal responses to defined nutrient deficiencies and can be combined with various tools, including genetics, to uncover metabolic pathways involved in nutritional adaptation. In addition, gut microbiota represents an additional layer that affects host nutritional environment. In many animals including *Drosophila*, commensal microbes have been increasingly recognized as key modulators of nutritional adaptation, although their involvement in specific micronutrient deficiencies remains largely unknown(*18*, *19*). In the case of *Drosophila* reared in laboratory conditions, its gut microbiota is primarily dominated by two families, *Acetobacteraceae* and *Lactobacillaceae*, both of which can influence host growth and metabolism under nutrient-limited conditions (*8*, *20–23*). In a pioneering study, Consuegra *et al.* combined microbiota manipulation with a chemically defined diet and showed that bacterial colonization can affect the survival of flies on a diet deficient in single nutrients (*8*). In the case of choline deficient diet, germ-free flies showed developmental arrest whereas *Acetobacter pomorum*, but not *Lactoplantibacillus plantarum*, colonization restored larval growth into pupae. These findings suggested that *A. pomorum* can buffer dietary choline deficiency, although the molecular mechanism remained unclear. More broadly, such results strengthen the view of *Drosophila* as a holobiont, whose phenotype emerges from interactions between the host genome and the microbial genes constituting its hologenome.

In this study, *Drosophila* genetics is paired with nutritional and commensal bacterial manipulation to uncover the mechanisms in place that prevent the flies from choline deficiency. We first show how a choline metabolite synthesized by a commensal bacteria can be utilized by the host as a nutrient source. We also demonstrate a mechanism by the host that functions independently of bacterial nutrient provisioning, whereby the host synthesize a choline substitute, β-methylcholine. This metabolite can be converted into βmPC and used in place of PC under dietary choline deficiency. Together, these findings reveal two independent metabolic pathways in the *Drosophila* hologenome that enable the host to overcome dietary choline deficiency.

## Results

### The effect of a choline-deficient diet depends on the presence of a microbiota

To investigate the organismal adaptive response to choline deficiency, we first examined its effects on *Drosophila* larval growth. We reared bleach-axenified embryos on either a control or choline-deficient diet (Fig. 1A). As expected, germ-free flies failed to pupariate under choline-deficient conditions and exhibited significantly reduced body weight at day 6 (Fig. 1B–D), confirming that choline is essential for larval growth.

**Fig. 1.**
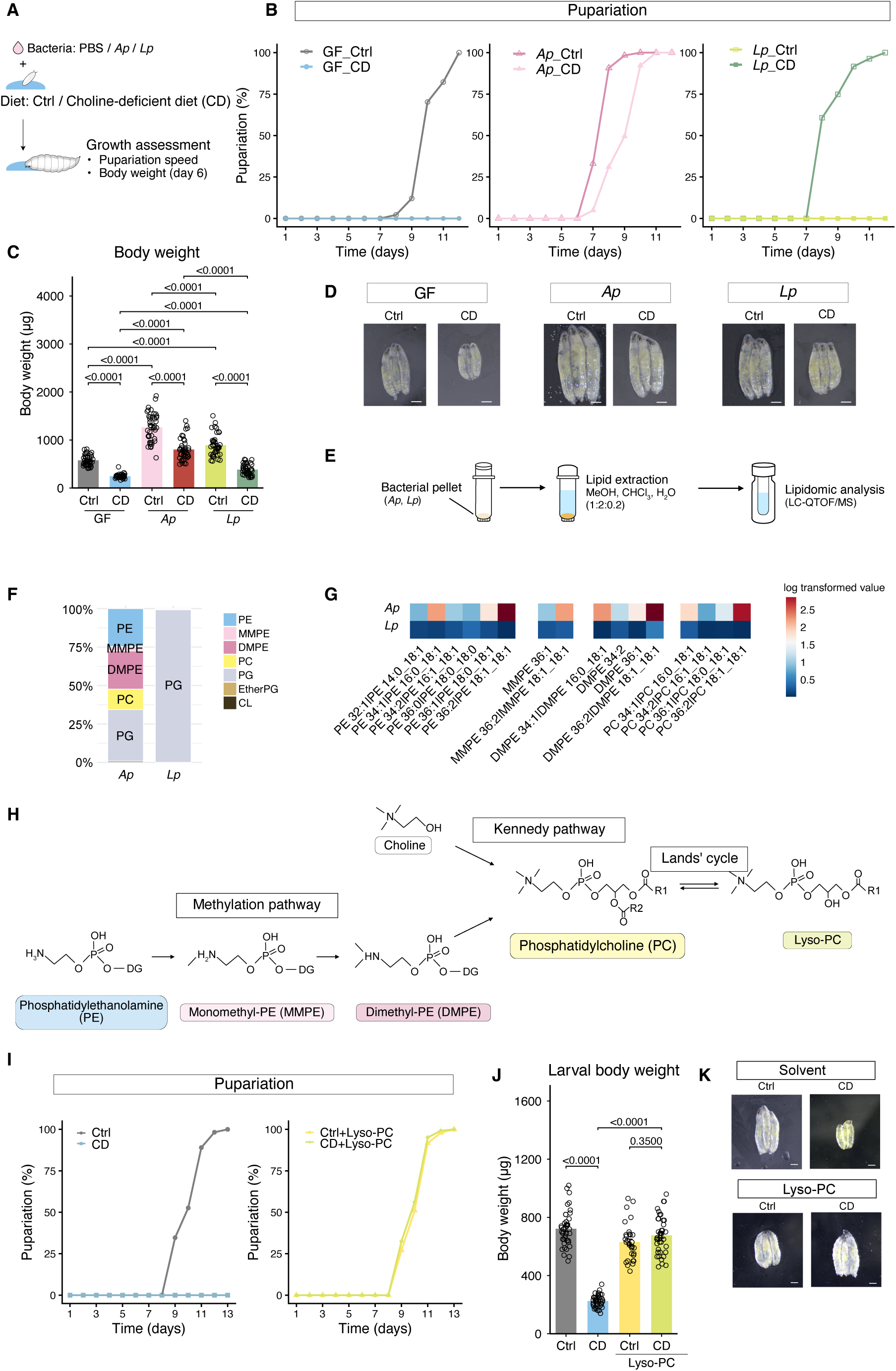
*Ap* supplementation rescues dietary choline deficiency by supplementing choline metabolites to the host. (**A**) Experimental scheme. GF embryos were plated to sterile control diet (Ctrl) or choline-deficient diet (CD), inoculated with control (PBS), Ap, or Lp. (**B**) Pupariation rates of larvae associated with *Ap* or *Lp* on CD; *n* = 168 (GF_Ctrl), *n* = 150-300 (GF_CD), *n* = 153 (Ap_Ctrl), *n* = 103 (Ap_CD), *n* = 191 (Lp_Ctrl), *n* = 150-300 (Lp_CD) (**C**) Bodyweight at day 6 of larvae associated with *Ap* or *Lp* on CD; *n* = 40. (**D**) Representative image at day 6 of larvae associated with *Ap* or *Lp* on CD. Scale bar = 0.5 mm. (**E**) Experimental scheme for lipidomic analysis on the bacteria. (**F**) Barplot showing phospholipid composition of *Ap* and *Lp*. (**G**) Heatmap showing the log 10 transformed concentrations of major phospholipids; *n* = 1. (**H**) Schematics of PC metabolism. PC can be synthesized by the Kennedy pathway which uses choline as the precursor, or the methylation pathway which uses PE as the precursor. In Land’ cycle, phosphatidylcholine is deacylated to lyso-PC and converted back to PC. (**I**) Pupariation rates of GF larvae on CD supplemented with1 mM lyso-PC ; *n* = 173 (Ctrl), *n* = 150-300 (CD), *n* = 96 (Ctrl+Lyso-PC), *n* = 123 (CD+Lyso-PC). (**J**) Body weight at day 6 for GF larvae on CD supplemented with1 mM lyso-PC; *n* = 40 (Ctrl), *n* = 31 (CD), *n* = 40 (Ctrl+Lyso-PC), *n* = 40 (CD+Lyso-PC) (**K**) Representative image at day 6 for GF larvae on CD supplemented with1 mM lyso-PC. Scale bar = 0.5 mm. CD, choline deficient diet; GF, germ-free; *Ap*, *Acetobacter persici* Ai; *Lp*, *Lactiplantibacillus plantarum* Lsi; PE, phosphatidylethanolamine; MMPE, monomethylphosphatidylethanolamine; DMPE, dimethylphosphatidylethanolamine; PC, phosphatidylcholine; PG, phosphatidylglycerol; CL, cardiolipin. Data points indicate biological replicate. For (**C** and **J**), statistical analysis was performed by one way analysis of variance (ANOVA). The *p* values were determined by post hoc analysis with Šídák’s multiple comparison test. All data are presented as mean ± s.e.m.

We have previously reported that the gut bacteria of flies in our laboratory condition was mainly composed of *Acetobacter persici* (*Ap*) and *Lactiplantibacillus plantarum* (*Lp*) (*24*). To determine which gut bacterial species can influence choline deficiency, we inoculated germ free embryos with bacterial isolates of *Ap* or *Lp*, and reared them on either control or choline-deficient diet (CD) (Fig. 1A). Under choline deficiency, both negative control (GF) flies and *Lp* inoculated flies failed to pupariate although body weight of *Lp* flies was larger than their germ-free counterparts (Fig. 1B-D). In contrast, *Ap* colonization fully rescued the pupariation rates, and increased larval body weight compared to GF or *Lp*-associated flies on choline deficient diet. The specific rescue of pupariation by *Ap* but not *Lp,* indicates the presence of an *Ap*-specific mechanism that enables the host to withstand choline deficiency. In contrast, both *Ap* and *Lp* is known to promote larval growth via various signalling pathways, which likely contributed to the increased body weight in *Ap* and *Lp* associated larvae independent of dietary choline (*20*, *21*, *23*, *25*). Nevertheless, even *Ap*-associated flies exhibited delayed pupariation and smaller body weight at day 6 on choline deficient diet relative to control diet, indicating that bacterial supplementation cannot fully compensate for the absence of dietary choline (Fig. 1B-D).

### *Ap*-derived phosphatidylcholine rescues choline deficiency

Given that commensal bacteria can provide nutrients to the host (*8*), we hypothesized that *Ap* may mitigate the effects of a choline-deficient diet by supplying choline or its metabolites to the host. In animals, choline is primarily metabolized into acetylcholine and phosphatidylcholine (PC). However, our previous LC–MS/MS analyses indicated that *Ap* did not synthesize choline or acetylcholine when cultured in MRS medium or in a standard fly food (*26*, *27*).

To determine whether *Ap* can synthesize PC, or related phospholipids, we conducted lipidomic profiling of *Ap* and *Lp*. Both bacterial isolates were cultured in MRS medium under standard aerobic conditions and subjected to LC–MS/MS analysis (Fig. 1E). The lipid profiles of *Ap* and *Lp* differed markedly (fig. S1). For instance, isoprenoids such as CoQ9 and CoQ10, along with cardiolipins, were abundant in *Ap* relative to *Lp*. This enrichment is consistent with the fact that *Ap* is an aerobic bacterium that relies on oxidative respiration for energy production. Conversely, glycolipids such as digalactosyldiacylglycerol (DGDG) were more prominent in *Lp*, which is typical for Gram-positive bacteria, where DGDG are known to serves as a membrane lipid anchor.

There were also distinct differences between the two bacterial species in their phospholipid compositions. While phosphatidylglycerol (PG) was the dominant phospholipid in both strains, it accounted for less than half of the total phospholipid content in *Ap* (Fig. 1F). The remaining fraction of *Ap*’s phospholipids consisted of PC, PE, monomethyl-PE (MMPE), and dimethyl-PE (DMPE), all of which were absent in *Lp* (Fig. 1F–G). PE, MMPE, and DMPE are intermediates in the methylation pathway which synthesizes PC from PE (Fig. 1H). These findings suggested that *Ap* may possess a functional methylation pathway. PC biosynthesis via methylation has previously been documented in other acetic acid–producing bacteria, including species within the *Acetobacter* and *Gluconobacter* genera (*28–30*).

In *Drosophila*, PC is predominantly synthesized from free choline via Kennedy pathway, and the existence of the methylation pathway has never been reported (Fig. 1H). Since the methylation pathway can function independently of environmental choline, we hypothesized that *Ap*, but not *Drosophila,* might be able to synthesize PC and transfer it to the host, regardless of choline content in the host diet. In insects, PC is thought to be absorbed as lyso-PC or choline since bioavailability of PC is very low (Fig. 1H) (*31*, *32*). As we expected, lyso-PC supplementation fully rescued both the pupariation defect and the larval body weight of germ-free animals observed under choline-deficient conditions (Fig. 1I–K). These results suggest that PC from *Ap* can be utilized by the host to mitigate choline deficiency.

### Bacteria-derived *PmtA* gene rescues choline deficiency

To identify the genes involved in PC biosynthesis by *Ap*, we analyzed the genome sequence of *Ap* and *Lp* that we have previously reported (*24*). We found that *Ap* has a single enzyme encoded by a gene, *Phosphatidylethanolamine methyltransferase A* (*PmtA*), that catalyzes all three reactions in the methylation pathway (Fig. 1H, Fig. 2A)(*33*). To determine whether *PmtA* alone is sufficient to rescue the effects of dietary choline deficiency, we generated transgenic flies carrying *ApPmtA* under the control of a UAS. Strikingly, overexpression of this bacterial gene in flies fully restored body weight and pupariation rates in larvae reared on a choline-deficient diet (Fig. 2B-C). This result demonstrated that PC biosynthesis via the bacterial *PmtA* gene is sufficient to ameliorate the effects of choline-deficient diet.

**Fig. 2.**
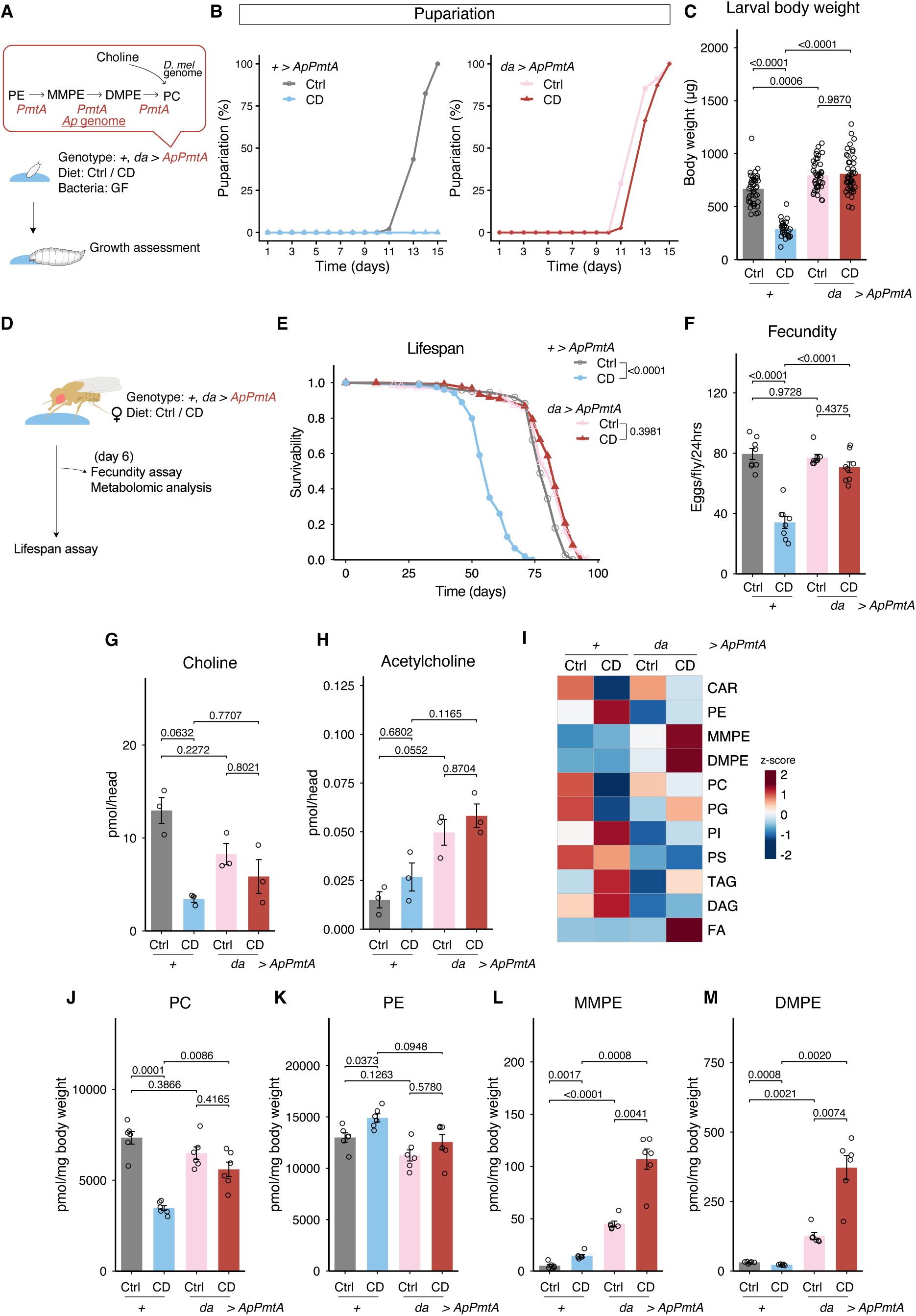
*Ap* derived *PmtA* gene overexpression rescues the effects of choline deficient diet. (**A**) Experimental scheme. *PmtA* gene derived from *Ap* genome was inserted to the fly genome under UAS. *PmtA* was overexpressed in the whole body using *da-gal4*. The GF flies were plated to CD and assessed for growth. (**B**) Pupariation rates of *da>ApPmtA* larvae on CD; *n* = 229 (Ctrl_*+>ApPmtA*), *n* = 150-300 (CD_*+>ApPmtA*), *n* = 193 (Ctrl_*da>ApPmtA*), *n* = 194 (CD_*da>ApPmtA*). (**C**) Bodyweight of day 6 *da>ApPmtA* larvae on CD; *n* = 29 (Ctrl_ +>*ApPmtA*), *n* = 40 (CD_*+>ApPmtA*), *n* = 40 (Ctrl_*da>ApPmtA*), *n* = 45 (CD_*da>ApPmtA*). (**D**) Experimental scheme for adult experiments. Females flies were sorted and placed on CD at day2. (**E**) Survival curve of *da*>*ApPmtA* flies on lifelong CD. *n* = 145 (Ctrl, +>*ApPmtA*), *n* = 154 (CD_*+>ApPmtA*), *n* = 139 (Ctrl_*da>ApPmtA*), *n* = 128 (CD_*da>ApPmtA*). (**F**) Results of egglaying assay for *da*>*ApPmtA* flies under CD for 6 days; *n* = 8. (**G-H**) Quantification of choline (**G**) and acetylcholine (**H**) in the head of *da>ApPmtA* flies after 6 days on CD; *n* = 3. (**I**) Lipid species that changed under CD and *da>ApPmtA*. The average concentrations of six samples were calculated, and the corresponding *z*-scores were visualized in the heatmap. (**J-M**) Quantification of PC (**J**), PE (**K**), MMPE (**L**), DMPE (**M**) in whole body of *da>ApPmtA* female flies on CD. CD, choline deficient diet; GF, germ-free; CAR, long-chain acyl-carnitine; PE, phosphatidylethanolamine; MMPE, monomethylphosphatidylethanolamine; DMPE, dimethylphosphatidylethanolamine; PC, phosphatidylcholine; PG, phosphatidylglycerol; PI, phosphatidylinositol; PS, phosphatidylserine; TAG, triacylglycerol; DAG, diacylglycerol; FA, fatty acid. For (**C**, **F-H, J-M**), statistical analysis was performed by one way analysis of variance (ANOVA). The *p* values were determined by post hoc analysis with Šídák’s multiple comparison test. For (**E**), statistical analysis was performed by log rank test with Holm correction. All data are presented as mean ± s.e.m.

### Choline is an essential nutrient in adult flies

Long-term feeding on a choline-deficient diet caused complete developmental arrest, making further analyses technically challenging. To explore whether similar compensatory mechanisms function in adults, we examined the effects of choline deficiency in adult flies. Adult flies were fed a choline-deficient diet beginning at day 2 post-eclosion (fig. S2A). Under these conditions, female lifespan was significantly shortened compared to males(fig. S2B). This sex-specific sensitivity may reflect the reliance of females on dietary choline for egg production, making them more vulnerable to its depletion(*34*). In addition, we observed that females maintained on a choline-deficient diet exhibited reduced ovary size and fecundity (fig. S2C-E), which is a typical hallmark of nutritional deficiency. Resistance to nutrient deprivation was also increased upon choline deficiency, suggesting altered energy homeostasis (fig. S2F). Morphological examination revealed that the fat body appeared visibly enlarged under choline-deficient conditions. LipidTOX staining showed that choline deprivation promotes the accumulation of larger lipid droplets in the abdominal fat body (fig. S2G, H). These data suggested that choline deficiency disrupts lipid homeostasis, similarly to mammalian liver.

Transcriptomic analysis revealed marked alterations in gene expression in female abdomen after 6 days of choline deficient diet (fig. S3A). We identified that 85 genes were upregulated, and 56 were downregulated (fig. S3B). Gene ontology analysis confirmed that genes involved in embryogenesis were particularly affected, consistent with our observation that choline is an essential nutrient for maintaining female reproduction (fig. S3C). Genes involved in lipid metabolism were also altered by choline deficiency (fig. S3D,E). For example, among the genes downregulated by choline deficiency was *1-Acylglycerol-3-phosphate O-acyltransferase 4* (*Agpat4*), which is involved in the synthesis of phosphatidic acid and diacylglycerol from glycerol-3-phosphate. *Lipid storage droplet-1* (*Lsd1)*, which promotes lipolysis by recruiting lipases to lipid droplets, was upregulated by choline deficiency. Similarly, *CG10814*, a gene in the carnitine biosynthesis pathway, was upregulated. Carnitine plays a central role in lipid metabolism by transporting fatty acids into mitochondria for β-oxidation. *CG10814* encodes γ-butyrobetaine dioxygenase, which catalyzes the final step of carnitine biosynthesis (fig. S3F)(*35*). We found that most genes in this pathway tended to be transcriptionally upregulated (fig. S3G). Taken together, these findings indicate that lipid homeostasis is aberrated under choline deficiency.

### *PmtA* can rescue the effects of choline deficient diet in adult female flies

To determine whether *Ap* can alleviate the effects of choline deficiency in adults, as observed in larvae, we added cultured *Ap* to the choline-deficient diet (fig. S4A). Contrary to our expectation, *Ap* supplementation did not restore egg-laying capacity in females (fig. S4B). Similarly, dietary supplementation with lyso-PC failed to fully rescue the reproductive defects caused by choline deficiency (fig. S4C). lyso-PC supplementation reduced starvation resistance in female flies under both control and choline deficiency, implying that lyso-PC was bioavailable and active (fig. S4D). Cox proportional hazards analysis confirmed that lyso-PC significantly influenced starvation resistance and partially ameliorated the effects of choline deficiency (Table S1). Together, these findings suggest that, in contrast to larval stage, *Ap*-derived PC cannot effectively compensate for dietary choline deficiency in adults.

Interestingly, overexpression of the *ApPmtA* entirely rescued the lifespan shortening effects by dietary choline deficiency in adult flies (Fig. 2D,E, Table S2). Fecundity and ovary size reduction caused by choline deficient diet was also restored completely (Fig. 2F, fig.S4E-F). In addition, overexpression of *ApPmtA* abolished the increased starvation resistance induced by dietary choline deficiency (fig. S4G, Table S3). These findings indicate that *PmtA* expression, rather than dietary supplementation, was sufficient to alleviate choline deficiency-induced phenotypes. This discrepancy could reflect (1) life stage–specific differences in phospholipid absorption/utilization, or (2) variation in bacterial physiology affecting phosphatidylcholine provisioning.

### *PmtA* rescues the effects of choline deficient diet on lipid metabolism

To quantify choline metabolites, we used LC-MS/MS to analyze choline and acetylcholine levels. Head samples were analyzed to exclude the effect of diet-derived choline in the gut and minimize potential confounding effects from the ovaries. Under control diet conditions, *ApPmtA* overexpression tended to decrease free choline levels and increase acetylcholine levels (Fig. 2G-H), suggesting an overall alteration of choline metabolism. When compared across dietary conditions, control flies exhibited markedly reduced choline levels on choline-deficient diet, a decrease that was not observed in flies overexpressing *ApPmtA*. In contrast, acetylcholine levels remained largely unchanged across dietary conditions.

We next performed lipidomic profiling and quantified phospholipids and other major lipid species, including triacylglycerols (TAGs) and fatty acids, in whole-body samples (Fig. 2I). In mammalian cells, PC is the most abundant phospholipid, followed by PE, but the insect cell membranes are known to contain proportionally less PC and more PE(*47*). Consistently, the whole body of the female flies exhibited a high PE/PC ratio (approximately 2 in control genotype on choline containing diet) (Fig. 2J and K). This compositional difference likely reduces the reliance of insects on PC, and may weaken the evolutionary incentive to retain the methylation pathway for maintaining the PE/PC ratio, potentially explaining its loss in the fly genome.

Principal component analysis (PCA) revealed that dietary choline strongly separated groups in the control genotype along both PC1 and PC2 axes, whereas this separation was less pronounced in *ApPmtA*-overexpressing flies, suggesting that *ApPmtA* expression partially rescued the metabolic effects of choline deficiency (fig. S5A-C). Lipid species contributing most to PC1 and PC2 included PE and its methylated derivatives, TAGs, fatty acids, and cardiolipins (fig. S5B-C). Total TAG and DAG levels tended to increase under choline-deficient conditions, possibly reflecting changes in general lipid metabolism or phospholipid biosynthesis (fig. S5D-E).

Most importantly, under choline-deficient conditions, PC levels decreased by approximately half, while PE levels increased; this alteration was ameliorated by *ApPmtA* overexpression (Fig. 2I–K, fig. S5F). Moreover, in *ApPmtA* overexpressing flies, the intermediate metabolites MMPE and DMPE were significantly elevated without a corresponding change in total PE levels (Fig. 2I, L, M, fig. S5F). Closer inspection of MMPE and DMPE revealed that PE species containing total acyl chain lengths of 30–34 carbons were preferentially utilized in the *ApPmtA*-mediated conversion to PC (fig. S5F). These results confirm that while there may be some bias in the acyl chain length of its substrate, the *ApPmtA*-dependent methylation pathway is functional *in vivo*. Interestingly, despite the additional PC biosynthesis pathway, *ApPmtA* overexpression flies did not show increased PC levels on control (choline-containing) diet, suggesting the presence of a robust mechanism that maintains internal phospholipid homeostasis. Collectively, these findings demonstrate that PC synthesized via *ApPmtA* is sufficient to prevent choline deficiency–induced developmental and physiological defects, underscoring the central role of PC biosynthesis in host–microbe metabolic interactions.

### Carnitine supplementation mitigates the effects of choline deficiency

To further characterize the effects of *ApPmtA*, we performed RNAseq analysis on abdominal tissues of *ApPmtA*-overexpressing flies on choline deficient diet (fig. S6A). *ApPmtA* overexpression rescued the transcriptional profile of 260 genes, including that of *CG10814*, which encodes the enzyme in the carnitine biosynthesis pathway (fig. S6B). Notably, upregulation of genes involved in carnitine biosynthesis under choline deficiency was suppressed under *ApPmtA* overexpression (fig. S6C), further corroborating that *ApPmtA* alleviates the transcriptional effects of choline deficiency.

In cells, carnitine exists as either free carnitine or acylcarnitine, its acylated derivatives. Using targeted metabolomics, we detected free carnitine and acetylcarnitine, while non-targeted lipidomic analysis allowed quantification of acylcarnitines with fatty acyl chains longer than 14 carbons due to technical constraints. Quantification showed that in control flies, carnitine, acetylcarnitine and long-chain acylcarnitine levels were drastically decreased under choline deficiency (fig. S6D-F). *ApPmtA* overexpression increased carnitine and acetylcarnitine levels under choline deficiency, with a similar, albeit non-significant, trend observed under the control diet. This suggests that *ApPmtA* overexpression elevates overall carnitine and acetylcarnitine abundance, but choline deficiency still imposes a general depletion regardless of genotype. In contrast, long-chain acylcarnitine levels were largely unaffected by *ApPmtA* overexpression in control diet. The decline observed under choline-deficient conditions was at least partially rescued as well, suggesting that long-chain acylcarnitine level maintenance is prioritized over free or acetylcarnitine in this condition. These results indicate that choline deficiency depletes total carnitine pools and that *ApPmtA* overexpression can partially restore their levels. Together with the transcriptomic data, these results imply that carnitine utilization is enhanced under choline deficiency, while carnitine biosynthesis is transcriptionally upregulated most likely as a response to depleted internal stores.

To directly investigate the relationship between dietary choline deficiency and carnitine depletion, we supplemented a choline-deficient diet with carnitine. Surprisingly, carnitine supplementation restored pupariation and larval body size (Fig. 3A–C). In adult females, carnitine alleviated the lifespan shortening caused by choline deficiency (Fig. 3D–E, Table S4). Notably, however, carnitine supplementation reduced lifespan on a control diet, suggesting that the high dosage may have exerted pathogenic effects. In contrast, reduction in fecundity by choline deficiency was completely restored (Fig. 3F), indicating that the effects of choline deficiency can be rescued by carnitine in adult flies as well as in larvae. Quantification of carnitine and its related metabolites by LC–MS/MS revealed that, although dietary supplementation greatly increased carnitine content, depleting choline from the supplemented diet still significantly reduced its levels (Fig. 3G-J). This suggests that carnitine utilization, and not its absolute abundance is key to rescuing the phenotypes. These data also indicate that flies possess a high capacity for carnitine consumption, especially under choline deficiency.

**Fig. 3.**
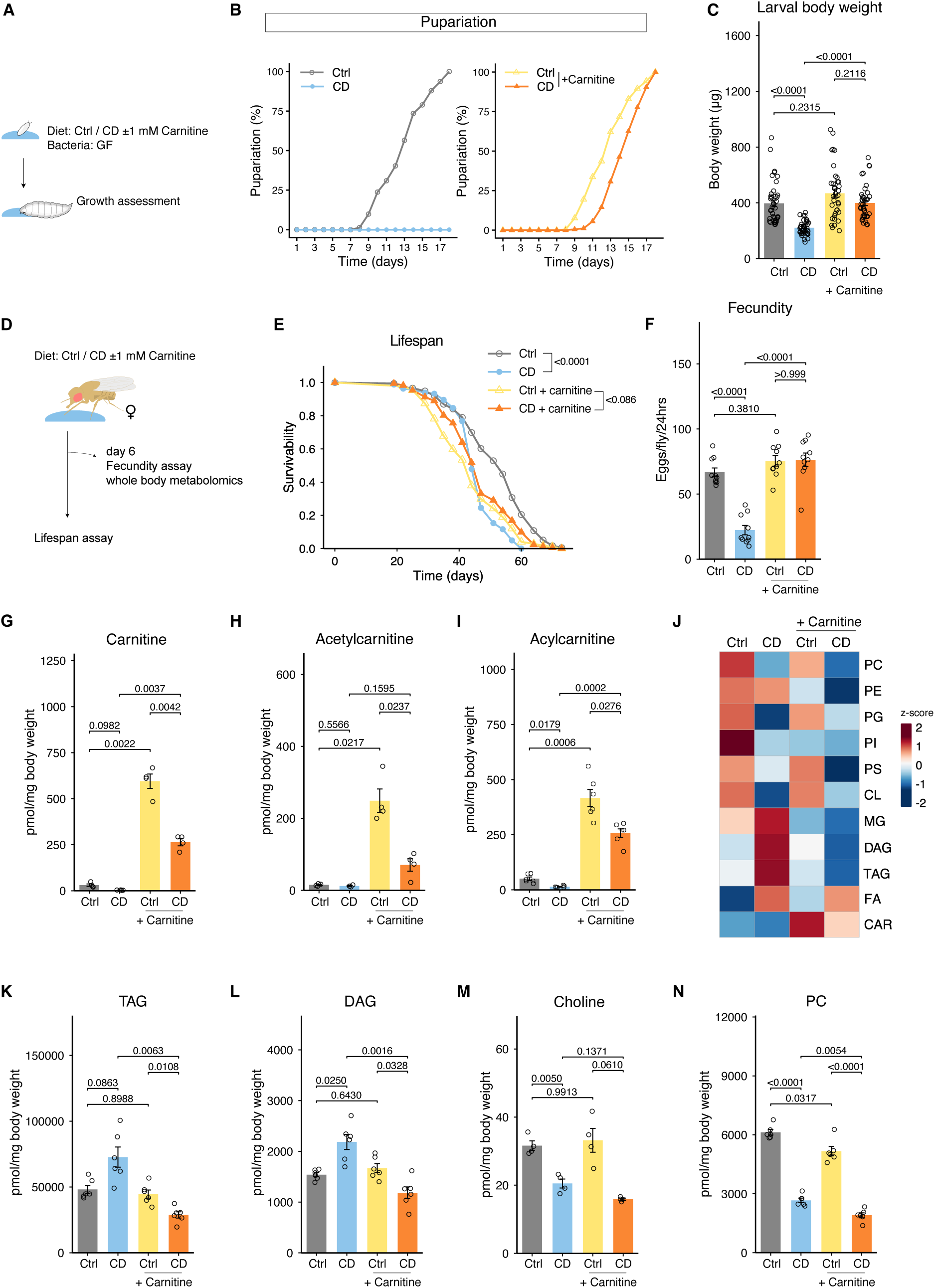
Carnitine supplementation to the diet can rescue the effects of choline deficiency, but not the deficiency of choline or its metabolites. (**A**) Experimental scheme. GF embryos were plated to sterile CD supplemented with 1 mM carnitine and assessed for growth. (**B**) Pupariation rates of GF larvae on CD supplemented with carnitine; *n* = 223 (Ctrl), *n* = 150-300 (CD), *n* = 211 (Ctrl + Carnitine), *n* = 179 (CD + Carnitine). (**C**) Bodyweight at day 6 of GF larvae on CD supplemented with carnitine; *n* = 40. (**D**) Experimental scheme. Female Canton-S flies were fed CD supplemented with 1 mM carnitine and assessed for lifespan and fecundity. (**E**) Survival curve for Canton-S female flies on lifelong CD supplemented with carnitine; *n* = 176 (Ctrl), *n* = 163 (CD), *n* = 174 (Ctrl + Carnitine), *n* = 172 (CD + Carnitine). (**F**) Fecundity of flies after 6 days on CD supplemented with carnitine; *n* = 10. (**G-I**) Quantification of carnitine (**G**), acetylcarnitine (**H**) and long-chain acylcarnitine (**I**) for whole body of flies after 6 days of CD supplemented with carnitine. (**J**) Lipid species that changed under CD or carnitine supplementation. The average concentrations of six samples were calculated, and the corresponding *z*-scores were visualized in the heatmap. (**K-N**) Quantification of TAG (**K**), DAG (**L**), choline (**M**), and PC (**N**) in the whole body of flies after 6 days on CD. CD, choline deficient diet; GF, germ-free. For (**C**, **F-I**, and **K-N**), statistical analysis were performed by one way analysis of variance (ANOVA). The *p* values were determined by post hoc analysis with Šídák’s multiple comparison test. For (**E**), statistical analysis was performed by log rank test with Holm correction. All data are presented as mean ± s.e.m.

We next examined the lipid profiles of carnitine-supplemented flies (Fig. 3J). Principal component analysis (PCA) revealed that control and choline-deficient groups were clearly separated along both PC1 and PC2 axes, whereas this separation along PC1 was abolished by carnitine supplementation (fig. S7A). Lipid species contributing most to PC1 included triacylglycerols (TAGs) and diacylglycerols (DAGs) (fig. S7B). Total TAG levels tended to increase and DAG levels rose markedly under choline deficiency, but these changes were absent upon carnitine supplementation (Fig. 3K–L). These findings indicate that sufficient carnitine availability can buffer the metabolic consequences of choline deficiency.

We first hypothesized that carnitine might be converted to choline or PC. However, no direct metabolic pathway linking carnitine and choline has been described to date. Consistent with this, carnitine supplementation did not restore internal choline or phosphatidylcholine (PC) levels under dietary choline deficiency (Fig. 3M-N, fig. S7C). In fact, carnitine supplementation, regardless of dietary choline content, modestly decreased PC levels. Collectively, these findings demonstrate that carnitine supplementation mitigates both the phenotypic and metabolic effects of dietary choline deficiency without restoring total choline or PC levels.

### Carnitine is metabolized into a phospholipid and rescued choline deficiency

In 1960s, carnitine has been serendipitously shown to functionally substitute choline, likely owing to their structural similarity, as both metabolites contain a trimethylamine residue (*36–38*). Theoretically, decarboxylation of carnitine can yield β-methylated choline (*36*), which was tentatively proposed to be able to be incorporated into a phospholipid named β-methylphosphatidylcholine (βmPC) (Fig. 4A). The presence of βmPC was proposed in 1961 where Bieber *et al.* fed *phormia regina* larvae with choline-deficient, carnitine-supplemented diet (*38*). They used paper chromatography to infer that the phospholipid contained β-methylcholine as its head group. Similar methods, namely paper chromatography or thin layer chromatography, were used for sample from other insects(*37*) as well as rats(*36*), most of which reported comparable findings. However, βmPC has never been directly identified by mass spectrometry.

**Fig. 4.**
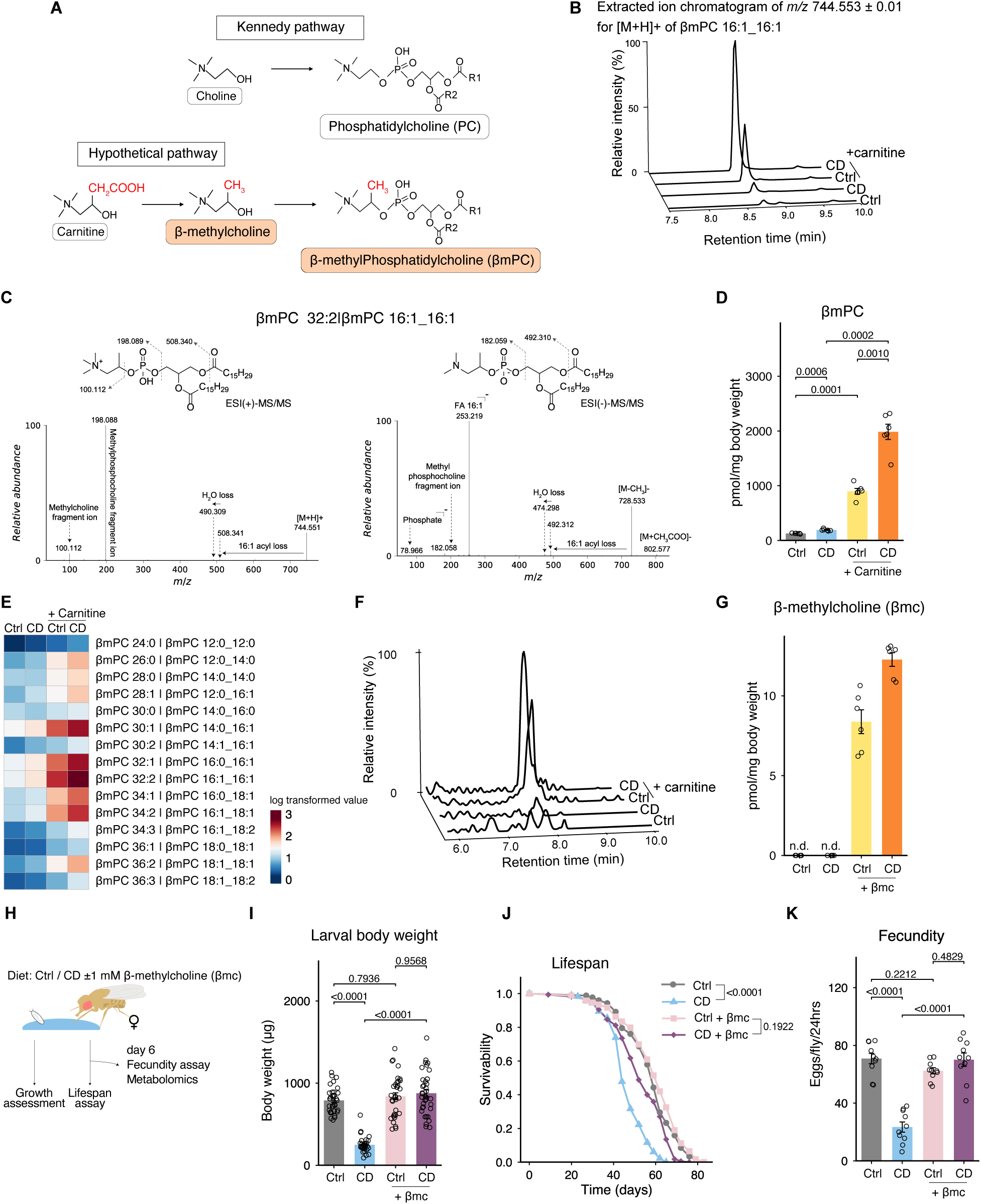
Carnitine can rescue choline deficiency because it is metabolized into β-methylcholine. (**A**) Hypothetical metabolic pathway for βmPC biosynthesis. Carnitine is decarboxylated to β-methylcholine, which is then metabolized into a phospholipid. Metabolite not previously identified as endogenous using LC-MS/MS is shaded in orange. (**B**) Chromatogram of methylated phosphatidylcholine named βmPC. Representative data from each dietary conditions for metabolite with 744.553 m/z are shown. (**C**) Representative image showing product ion for βmPC with 744.553 m/z for both positive and negative ion mode. (**D**) Quantification of βmPC. n.d. implies not detected. ; *n* = 6. (**E**) All species of annotated βmPC. The average concentrations from six samples were calculated, and the corresponding *z*-scores were visualized in the heatmap. (**F**) Mass chromatogram of β-methylcholine (βmc) at m/z 119.00>61.05. (**G**) Quantification of βmc; *n* = 6. (**H**) Experimental schematics. GF embryos or adult female flies were placed on control diet (Ctrl) or choline-deficient diet (CD) containing 1 mM βmc. (**I**) Body weight of day 7 larvae on CD supplemented with βmc; *n* = 37 (Ctrl), *n* = 36 (CD), *n* = 36 (Ctrl+βmc), *n* = 39 (CD+βmc). (**J**) Survival curve for adult female flies on lifelong CD supplemented with βmc; *n* = 148 (Ctrl), *n* = 137 (CD), *n* = 180 (Ctrl + βmc), *n* = 167 (CD + βmc). Data points indicate biological replicate. (**K**) Fecundity of flies after 6 days on CD supplemented with βmc; *n* = 10. CD, choline deficient diet; GF, germ-free. For (**D**, **I** and **K**), statistical analysis was performed by one way analysis of variance (ANOVA). The *p* values were determined by post hoc analysis with Šídák’s multiple comparison test. For (**J**), statistical analysis was performed by Log rank test with holm correction. All data are presented as mean ± s.e.m.

We attempted to detect βmPC using LC-MS/MS (Fig. 4B-D). Since we speculated that βmPC would have similar retention time to PC, we focused on phospholipids that were falsely annotated as PC. Indeed, we were able to detect a lipid species that were falsely identified as PC, which had a distinct peak at *m/z* of 100.068 and 198.112, which indicated the presence of an additional single methyl residue on the alpha or beta position of choline. The abundance of methylated PC species was influenced by both choline and carnitine availability (Fig. 4D). More specifically, a choline-deficient diet induced a slight increase in methylated PC relative to the control, while carnitine supplementation caused a more pronounced elevation regardless of dietary choline content. The acyl groups incorporated into βmPC was similar to that in PC, with total acyl chain lengths of 30, 32, and 34 carbons being most abundant (Fig. 4E). These results suggest that methylated PC is synthesized through the incorporation of decarboxylated carnitine into phospholipids, supporting the interpretation that this compound corresponds to βmPC.

Using LC–MS/MS, we also detected β-methylcholine, the head group of βmPC, in flies. β-methylcholine was readily detected in flies fed diets containing carnitine, whereas its levels in flies reared on diets without carnitine supplementation were below the detection threshold equivalent to approximately 2 pmol/mg body weight (Fig. 4F-G). Taken together, the results suggest that carnitine is converted to β-methylcholine, which act as a precursor to produce βmPC.

We next analyzed βmPC levels in *ApPmtA*-overexpressing flies (fig. S8). Consistent with the fluctuating carnitine and acetylcarnitine levels (Fig S6D-F), *ApPmtA* overexpression reduced βmPC content under both control and choline-deficient conditions. This result suggests that overall PC abundance, or at least its synthesis via the methylation pathway, influences βmPC production. Notably, choline deficiency increased βmPC levels in the whole body regardless of genotype. Although total PC levels were largely maintained by *ApPmtA* overexpression even under choline-deficient conditions (Fig. 2J), whether PC levels are maintained uniformly across all tissues remains unclear. Therefore, local availability within specific tissues or cell types might be driving this upregulation, elucidating the sensitivity in the regulation of this biosynthesis pathway.

Finally, we tested whether the presumed intermediate metabolite, β-methylcholine, can functionally substitute for choline (Fig. 4H). β-methylcholine supplementation rescued the decreased body weight (Fig. 4I). In adult female flies, β-methylcholine rescued, at least partially, the lifespan shortening induced by choline deficiency (Fig. 4J ,Table S5), and decline in fecundity (Fig. 4K). Taken together, our results highlight the importance of β-methylcholine production from carnitine, in overcoming choline deficiency.

## Discussion

Our present study in *Drosophila* uncovered two independent metabolic pathways that reduce dependence on dietary choline by sustaining PC or PC-like phospholipid pools. Dietary requirement of choline varies depending on genetic, dietary, and environmental factors (*6*, *9*, *40*) showcasing the complex nature of nutritional availability. In 2000, Joshua Lederberg proposed viewing hosts and their symbionts as a single “superorganism”, integrating genomes from both hosts and symbionts into one functional entity (*41*). *ApPmtA*-overexpressing flies can be seen as a prototype of such superorganismal metabolic homeostasis with respect to choline metabolism. The methylation pathway of PC synthesis is evolutionarily conserved in most eukaryotes, from yeast to mammals (*12*, *15*). In mammals, *PEMT* is expressed primarily in the liver, where it contributes to PC synthesis. Interestingly, the *PEMT* gene is largely absent in arthropods, raising the hypothesis that flies might outsource PC biosynthesis to their microbial partners under conditions where dietary choline is scarce. In the wild, PC supply is generally abundant owing to the microbes. In addition to *Acetobacter*, flies in the wild are highly associated with yeast (*45*), some of which has also been shown to conserve the methylation pathway for PC synthesis (*46*).

Bacterial PC synthesis is relatively uncommon, with only about 15% of bacterial species predicted to possess this capacity, predominantly within the Proteobacteria (*48*). In acetic acid producing bacteria, PC has been reported to contribute to tolerating acidic conditions(*28*, *29*).

For other bacterial species, PC has been shown to promote symbiosis with eukaryotic hosts, largely in the context of virulence (*49–51*). Non-bacterial pathogens are likewise known to exploit PC to enhance infection (*52–54*). To our knowledge, this is the first demonstration that commensal gut bacteria produce PC which serves as a nutrient source for the host. Given that *Acetobacter* and related acetic acid–producing bacteria are widely associated with arthropods, this mechanism is likely conserved especially across insect lineages(*55*). Nutritional provisioning by the bacteria is an important factor for most host-bacteria symbiosis, not limited to insects, so such mechanisms may also be conserved in many other animals, including mammals. Taken together, our findings highlight that an animal’s nutritional requirements are not fixed but are dynamically shaped by its gut microbial community.

Previously, we have shown that *Acetobacter* association can shorten *Drosophila* lifespan, likely due to immune activation or altered purine metabolism, yet also promote growth, fecundity, and stress resistance, which we proposed are traits consistent with a “live fast, die young” strategy (*26*, *56*, *57*). In contrast, under choline-deficient conditions, we found that *A. persici* association or *ApPmtA* overexpression enhanced growth and fecundity and even extended lifespan. This context-dependent reversal suggests that the effects of symbiosis may shift from a trade-off to a net benefit under host nutrient scarcity.

In this study, we focused on carnitine because of its pronounced decrease and altered metabolism under choline-deficient conditions. Our data suggested that reduced carnitine levels may reflect not merely depletion but an adaptive mechanism in which carnitine is metabolized to produce molecules that can functionally substitute for choline, namely βmPC and β-methylcholine. We quantified these compounds using LC–MS/MS, for the first time, establishing a mechanistic link between choline scarcity and carnitine metabolism. Choline restriction leads to carnitine depletion also in mammals including rats, and this has been thought to be the result of aberrant lipid metabolism or indirect effects of altered one carbon metabolism(*40*, *58*). While some studies suggest the presence of βmPC in mammals, its presence remains elusive(*59*, *60*). The presumed enzyme was isolated only once to our knowledge by Khairallah and Wolf in 1967 in a study using rats (*36*), and the genes involved had never been identified. In addition, some studies have proposed that phosphatidylcarnitine, instead of free carnitine, is decarboxylated into β-methylphosphatidylcholine(*59*, *61*). Validation of the βmPC biosynthesis pathway, and its conservation across species require additional exploration.

Carnitine is an important metabolite for energy homeostasis due to its central role in maintaining mitochondrial function and facilitating lipid oxidation(*39*). Concordant with this, several studies have suggested that carnitine supplementation can mitigate the effects of lipid accumulation caused from metabolic diseases(*62*, *63*). In humans, carnitine and its derivatives are commercially available as nutritional supplements, and clinical trials in metabolically impaired individuals have reported beneficial metabolic outcomes (*64*, *65*). However, our data open the possibility of an additional mode of action for carnitine in addition to its canonical mitochondrial functions. Specifically, some of the positive effects of carnitine supplementation may arise from the synthesis of βmPC. Disentangling the physiological effects of carnitine from those of βmPC or β-methylcholine will be important for defining the underlying mechanisms and could guide more targeted therapeutic strategies.

## Materials and methods

### Drosophila husbandry

Flies were raised on a standard yeast-based diet containing 4.5% cornmeal (NIPPON CORPORATION), 6% brewer’s yeast (ASAHI BREWERIES, HB-P02), 6% glucose (Nihon Shokuhin Kako), and 0.8% agar (Ina Food Industry S-6) with 0.4% propionic acid (Wako 163-04726) and 0.15% butyl p-hydroxybenzoate (Wako 028-03685). Flies were maintained under conditions of 25°C. To synchronize the development, embryos were collected by agar plates (2.3% agar, 1% sucrose, and 0.35% acetic acid) with live yeast paste. Density was managed by collecting the embryo using PBS and plating a set volume of embryo per vial (12 µL per bottle of food). Adult flies were allowed to mate for 2 days after eclosion before being used in experiments. The fly lines used in this study were Canton-S, *w^Canton-S^*, *da-Gal4*(*66*), *Lsd2-YFP* (Kyoto DGGR, 115301) and *UAS-ApPmtA* (Generated in this study).

### Creation of *ApPmtA* expressing transgenic flies

To create transgenic flies expressing Ap-derived *PmtA* (*ApPmtA*) under a UAS promoter, we first performed codon optimization on the *PmtA* sequence and obtained the sequence shown below. *PmtA* sequence was acquired from the whole genome sequencing analysis result which we had previously reported(*24*). Codon optimization was performed by FASMAC (https://fasmac.co.jp/).

GAATTCCAACATGAGCGATGTCCTCCATGAATGCCCTCCAGGACAGGGCGAGGATCCAGTACCGCCTCCCAGCCGATCCGCGCTTGATGCAGAAGCAGTAAAGGTTGCGTATCGTAGGTGGGCTGCGGTTTACGACACGGTGTTCGGTGGCGTCAGCGGAGTTGGCAGGAAACGTGCAGTGGAGGCTGTGAACGCACTTCCCGGCACAAAAGTCCTGGAAGTTGGTGTCGGAACAGGCCTCGCACTGCCATGTTACACCCAAAACAAGAAGATCACCGGAATAGATTTGAGCGGCGATATGCTCGCTCGTGCTCGAGAGCGAGTCCAGAAGGAACACCTGACTAATGTAGAGGCGTTGCTTGAAATGGACGCGGAGGAGACTTCGTTCGCGGACGCAAGTTTCGATATAGCCGTGGCTATGTTTGTGGCCAGCGTCGTCCCGCATCCGCGTAAGCTGCTGAGTGAGCTGAAGAGGGTAGTGCGCCCGGGCGGACACATCCTGTTCGTGAATCACTTTCTGGCCCCAAACGGCGTCCGCGGCGCCGTTGAGCGCGGCATGGCTCGCGCCAGCCGCTCGCTGGGCTGGCATCCCGACTTTGCCATGGAGTCCCTGCTCCCGCCAGACGACCTGGCCCGCGCACAAATCCAGCCAGTTCCACCACTGGGCCTGTTTACCCTTGTTACTCTGGACCGCACTTAAGAATTC

The sequence above along with Kozak sequence was inserted into pUASt. The inserted plasmid was injected into *w1118* embryo by WellGenetics (https://wellgenetics.com/).

### Creation of Germ-free flies

Creation of germ free flies were done using the method previously described(*27*). The embryos were collected on an agar plate were rinsed in PBS and transferred to a small basket. The embryos were then washed with 500 mL of 70 % ethanol twice and dipped in a 3% sodium hypochlorite solution for 5 minutes. The embryos were then washed with 1 L tap water and plated to sterile food that had been UV sterilized for at least 20 minutes. The vials were kept in a sterile plastic container. The bacterial condition was checked by plating the food onto MRS agar plate right before the experiment. Results obtained from flies reared in contaminated vials were excluded post analysis.

### Generation of gnotobiotic flies

*A. persici* Ai and *L. plantarum* Lsi(*24*), was isolated previously from flies maintained in the laboratory. Glycerol stocks of the isolated bacteria were preserved in -80°C. The stocks were cultured in roughly 5 mL of MRS (Oxoid, CM0359) stock at 30°C. Bacterial proliferation was measured by checking its absorbance (OD600) using DEN-600 Photometer (Funakoshi, BS-050109-AAK). The bacteria were cultured until the absorbance (OD600) reached approximately 0.5.

To create a gnotobiotic environment, bacterial culture was centrifuged at 25°C, 8000 × g for 5 minutes and washed in PBS. The bacterial sediment was resuspended and diluted in PBS until OD600 was 0.01. 100 µL of the resuspended solution was inoculated to the food that had been UV sterilized for at least 20 minutes. For control food, 100 µL of PBS was plated. For creation of gnotobiotic flies, germ free embryos were plated to the food manually.

### Dietary manipulation

For dietary manipulation, we used a chemically defined diet, also called holidic diet (*66*). Control diets were made by supplementing choline solution to CD. For larval diet, composition was adjusted as previously reported(*66*). For use in gnotobiotic experiments, nipagin and propionic acid was excluded from the food.

Nutrients were supplemented to the diet before aliquoting to individual vials. For supplementation of lyso-PC to the diet, 100 mM stock was made by dissolving S-lysoPC (Echelon Biosciences, L-1518) in EtOH. For carnitine or β-methylcholine, 500 mM or 100 mM stock was made by dissolving L-Carnitine (TCl, C0049) or β-methylcholine (TCl, M1293) in ultrapure water, respectively. Same volume of solvent was supplemented to the diet for controls.

For *Ap* supplementation, the bacteria was cultured as mentioned above, then washed and resuspended in PBS to make a solution with OD600 of 1.0. 100 µL of bacteria solution was added to the surface of the holidic diet. PBS was added for controls. The vials were left in room temperature until dry.

### LC-MS/MS analysis of compounds below 10-kDa

Ultra high performance LC–MS/MS (LCMS-8050/LCMS-8060NX, Shimadzu) based on the Primary metabolites package v.2 (Shimadzu) was used to quantify small compounds. First, samples were collected then homogenized in 150 μl of 80% methanol containing 10 μM of internal standards (methionine sulfone and 2-morpholinoethanesulfonic acid). For head samples, 8 heads were dissected in PBS. For whole body samples, 3 flies were weighed before being collected.

The homogenized samples were deproteinized by mixing with 75 μl of acetonitrile then transferred into a 10-kDa centrifugal device (Pall, OD010C35). The flowthrough of the sample was completely evaporated using a centrifugal concentrator (TOMY, CC-105), then resolubilized in ultrapure water. The samples were then injected to the LC–MS/MS with a PFPP column (Discovery HS F5 (2.1 mm × 150 mm, 3 μm), Sigma-Aldrich) in the column oven at 40 °C. The samples were separated using a gradient from solvent A (0.1% formic acid, water) to solvent B (0.1% formic acid, acetonitrile) for 20 min. Optimization of multiple reaction monitoring (MRM) parameters were performed by injecting the standard solution, then performing peak integration and parameter optimization with a software (LabSolutions, Shimadzu).

### LC-MS/MS analysis of lipophilic compounds

For sample collection, 10 female flies were weighed then frozen in -80°C. Bacteria was cultured as described above, to acquire absorbance (OD600) of 1.0. 10 mL of the culture was centrifuged and washed with PBS as described, then frozen in -80°C.

Total lipids were extracted from flies and bacteria using a previously described single-phase extraction method(*67*). Briefly, 200 µL of methanol with 5 µL of EquiSPLASH (Avanti Research), 0.32 µL of 10 mM FA 16:0-d3 and FA18:0-d3 (Olbracht Serdary Research Laboratories) and 100 µL of chloroform were added to 1 mg (bacterial dry pellet) or 2.5 mg (fly whole body) samples and incubated at room temperature for 2 hours. Then, 20 µL of water was added and centrifuged at 1680 × *g* for 10 min. The supernatant was transferred to a glass vial. Untargeted lipidomics was performed as previously described(*68*). ACQUITY UPLC system (I-Class; Waters) was coupled to QTOF-MS (TripleTOF 6600; Sciex), and lipids were separated on an Acquity UPLC Peptide BEH C18 column (50 × 2.1 mm, 1.7 µm; Waters). The mobile phase consisted of (A) 1:1:3 (v/v/v) acetonitrile:methanol:water, and (B) 100% 2-propanol. The gradient method and MS parameters were same as those in the previous study(*68*). Both mobile phase solvents were supplemented with ammonium acetate (5 mM) and 10 nM EDTA. Data analysis was performed using MS-DIAL bootstrap version 5.1.231129(*69*). Lipids were quantified at LSI level 3 using internal standards of the same or similar lipid classes, or representative standard compounds (https://lipidomicstandards.org/). The lipidomics minimal reporting checklist can be found in the Supplementary materials (Data S1 and 2).

### Larval growth assay

To evaluate the growth speed of the flies, larval bodyweight and pupariation timing was assessed. To measure the larval bodyweight, larvae were first floated up using 30 % glycerol solution and lightly washed in Milli-Q water. The larvae were then gently wiped down with a tissue to get rid of excess water then measured using a scale (Mettler Toledo., XPR2)

To quantify the pupariation speed, the number of pupae was counted daily. The experiment was terminated as adults eclosed and died on the medium, to exclude the effects of such additional nutrient source. Pupariation rate for each timepoint was calculated by setting the total number of pupae at the end of experiment as 100%. Vials containing more than 60 pupae at the end of the experiment were deemed crowded and excluded from the analysis.

### Egg laying assay

The following egg laying assay was performed to measure the reproductive ability of the female flies. Female flies post treatment and male Canton-S flies were lightly anesthetized by CO_2_, then three female flies and two male flies were transferred to a new vial containing yeast-based diet. The number of eggs laid was counted 24 hours later.

### Starvation resistance assay

To measure resistance to starvation, adult flies were transferred to a vial containing 1% agar diet. The number of dead flies were counted 1-4 times a day and the survival rate was calculated for each time point.

### Lifespan assay

For lifespan analysis, the viability was recorded every 3 to 4 days when they were flipped to a new vial containing fresh food. Minimum of six vials were used per condition.

### Transcriptomic analysis

For RNA sequencing analysis, abdominal tissues were dissected in PBS. Gut, Malpighian tubules, and reproductive tissues were removed using forceps. Total RNA from 15 female abdominal tissues, three or four replicates per condition, were extracted using Relia Prep^TM^ RNA Miniprep Kit (Promega, Z6112). Library preparation was performed by the RIKEN BDR Technical Support Facility using Illumina stranded mRNA prep ligation kit (Illumina, 20040534) and IDT for Illumina RNA UD indexes set A/B Ligation kit (Illumina, 20040553/20040554).

PCR cycle number was determined using KAPA real-time library amplification kit (Roche, KK2702 ), and library quality was verified using TapeStation HS D1000 assay. The samples were then sent to AZENTA and sequenced by Illumina Hiseq X to obtain paired end 150 base pair sequence data.

The RNA-seq data were processed as follows: quality check of the data were performed by FastQC (v.0.12.1)(*70*) and MultiQC (v.1.14)(*71*). Adaptors and low-quality bases were removed using Trim Galore (v.0.6.10)(*72*). Alignment to the *Drosophila genome* (BDGP6.54.114) was performed by hisat2 (v.2.2.1), read count calculation were done by Stringtie (v.2.2.1)(*73*), and differentially expressed genes were identified by edgeR (v.4.4.2)(*74*). RNA sequencing data have been deposited at the DDBJ under accession number PRJDB39867 and PRJDB39859.

Gene ontology analysis was performed using the R package clusterProfiler, using the function enrichGO(*75*).

### Imaging analysis

For analysis of ovary size, ovary was dissected in PBS on a silicone pad. Images were captured by stereomicroscope (Leica Microsystems GmbH, MZ10F).

For staining of lipid droplets, adult abdominal tissue was dissected in PBS. The carcass along with fat body was pinned to a silicone pad using insect pin and fixed in 4% PFA for 20 minutes at room temperature. The samples were washed multiple times with PBST buffer (0.1% Triton X-100 with PBS) and blocked with 5% normal donkey serum in PBST for 1 hour at room temperature. The samples were then incubated overnight at 4 °C with anti-GFP (rat IgG2a) monoclonal antibody (Nacalai tesque, 04404-26, 1:1000 dilution). The samples were washed with PBST buffer and incubated at room temperature for two hours with Alexa Fluor Plus 488 donkey anti-rat IgG (Thermo Fisher, A48269, 1:500 dilution), Hoechst 33342 (diluted to 0.4 mM; Invitrogen, H3570, 1:100 dilution), and Lipidtox deep red neutral lipid stain (invitrogen, H34477, dilution 1:250). After washing, tissues were mounted in Fade Gold. The images were captured by a confocal microscope (Leica Microsystems GmbH, TCS SP8).

Ovary and lipid droplet areas were quantified using Fiji software(*76*). For lipid droplets, regions of interest (ROIs) were selected manually. The images were thresholded and converted to binary, after which the area was quantified using the ’Analyze Particles’ function. The selection of particles was manually adjusted as needed.

### Statistical analysis

Statistical analysis and data visualization was performed using R. Unpaired student’s t-test was used to compare two groups. One-way ANOVA with Šídák’s multiple comparison test was used to compare multiple groups. For lifespan analysis, log rank test was used. Log rank test with holm correction was utilized for multiple comparisons. To analyze the interactions between various factors, dietary and genotype, Cox proportional hazards (PH) analysis was performed using the survival package(*77*). Hazard ratio and statistical significance based on z tests are reported in supplementary tables. Biological, not technical, replicates are represented by data points. Bar graphs are drawn as the means and SEM.

## Acknowledgments

We thank the Kyoto Stock Center for Drosophila stocks. We thank all members of Obata’s lab and Arita’s lab for technical assistance and advise. We would also like to acknowledge Tadashi Uemura for invaluable advice.

## Funding

AMED PRIME 20gm6310011 (FO)

AMED Moonshot Research and Development Program JP22zf0127007 (MA)

AMED NeDDTrim 21ae0121036 (MA)

JST FOREST JPMJFR2337 (FO)

JST ERATO ARITA Lipidome Atlas Project JPMJER2101 (MA)

Naito Foundation (FO)

Yakult Bio-Science Foundation (FO)

The Japan Society for Promotion of Science 25KJ1633 (YF)

Author contributions:

Conceptualization: YF, MA, FO

Investigation: YF, SM

Formal Analysis: YF, SM

Visualization: YF, SM

Funding acquisition: MA, FO

Supervision: HK, MA, FO

Writing – original draft: YF, FO

Writing – review & editing: YF, SM, HK, MA, FO

## Competing interests

Authors declare that they have no competing interests.

## Data and materials availability

All raw LC-MS/MS data are available from the MB-POST (https://repository.massbank.jp/) via the index MPST000124. All raw RNA-seq data are available from DDBJ (https://www.ddbj.nig.ac.jp/index-e.html) via the accession code PRJDB39867 and PRJDB39859. Any materials or additional information can be requested from the corresponding authors.

## Notes

### Competing Interest Statement

The authors have declared no competing interest.

