## Supplementary Figures for "Dual Metabolic Compensation to Dietary Choline Deficiency by the *Drosophila* Hologenome"

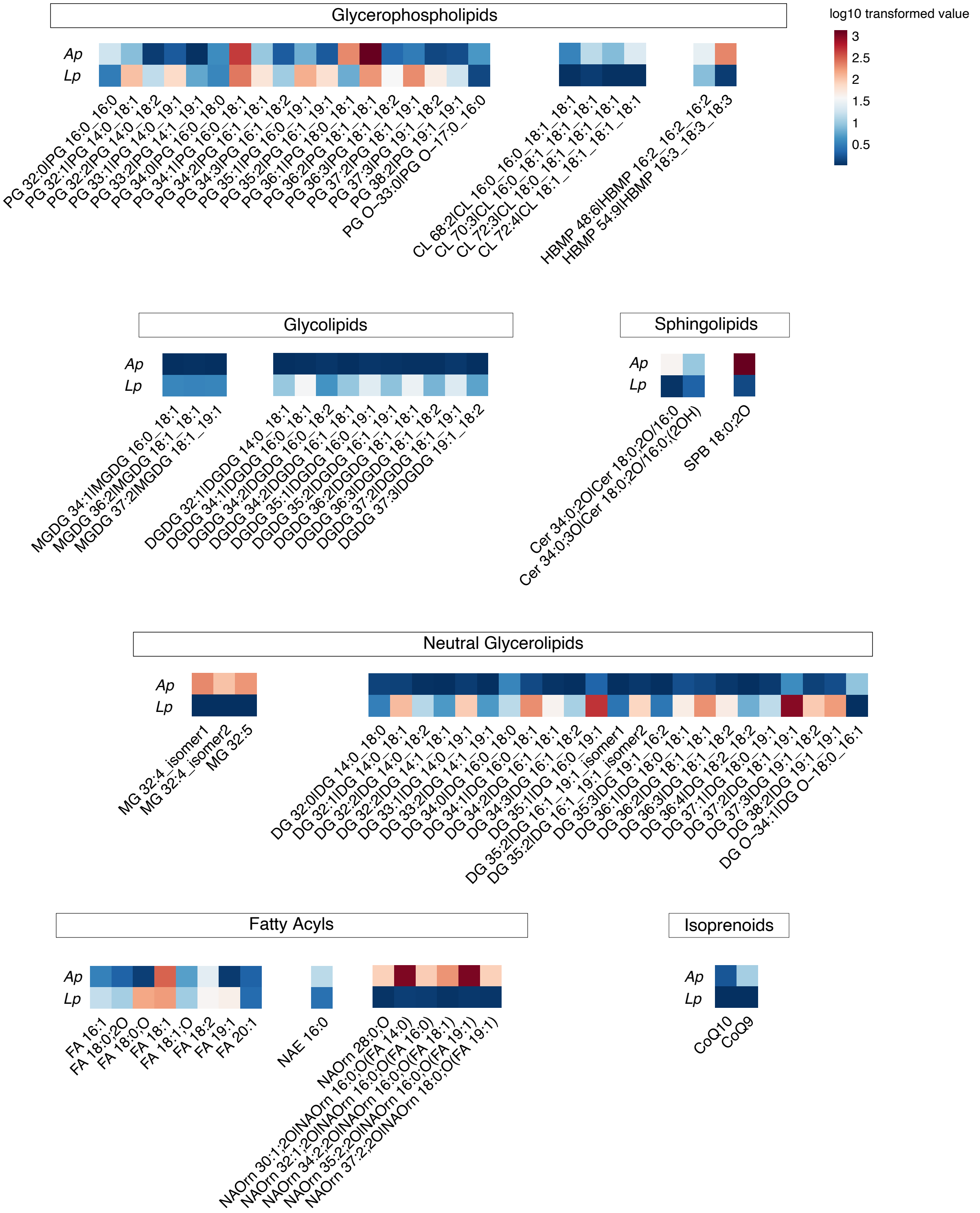


**Fig. S1. Lipid composition in *Drosophila* commensal bacteria**

Lipidome analysis of *Ap* and *Lp* pellets. Heatmap shows log 10 transformed concentrations of lipid species that were especially different between the two bacteria. *n* = 1. *Ap*, *Acetobacter persici* Ai; *Lp*, *Lactiplantibacillus plantarum* Lsi; PG, phosphatidylglycerol; CL, cardiolipin;

HBMP, hemibismonophosphatidic acid; MGDG, monogalactosyldiacylglycerol; DGDG, digalactosyldiacylglycerol; Cer, ceramide; SPB, sphingoid base; DG, diacylglycerol; MG, monoacylglycerol; FA, fatty acid; NAE, N-acylethanolamine; NAOrn, N-acylornithine; CoQ, coenzyme Q.

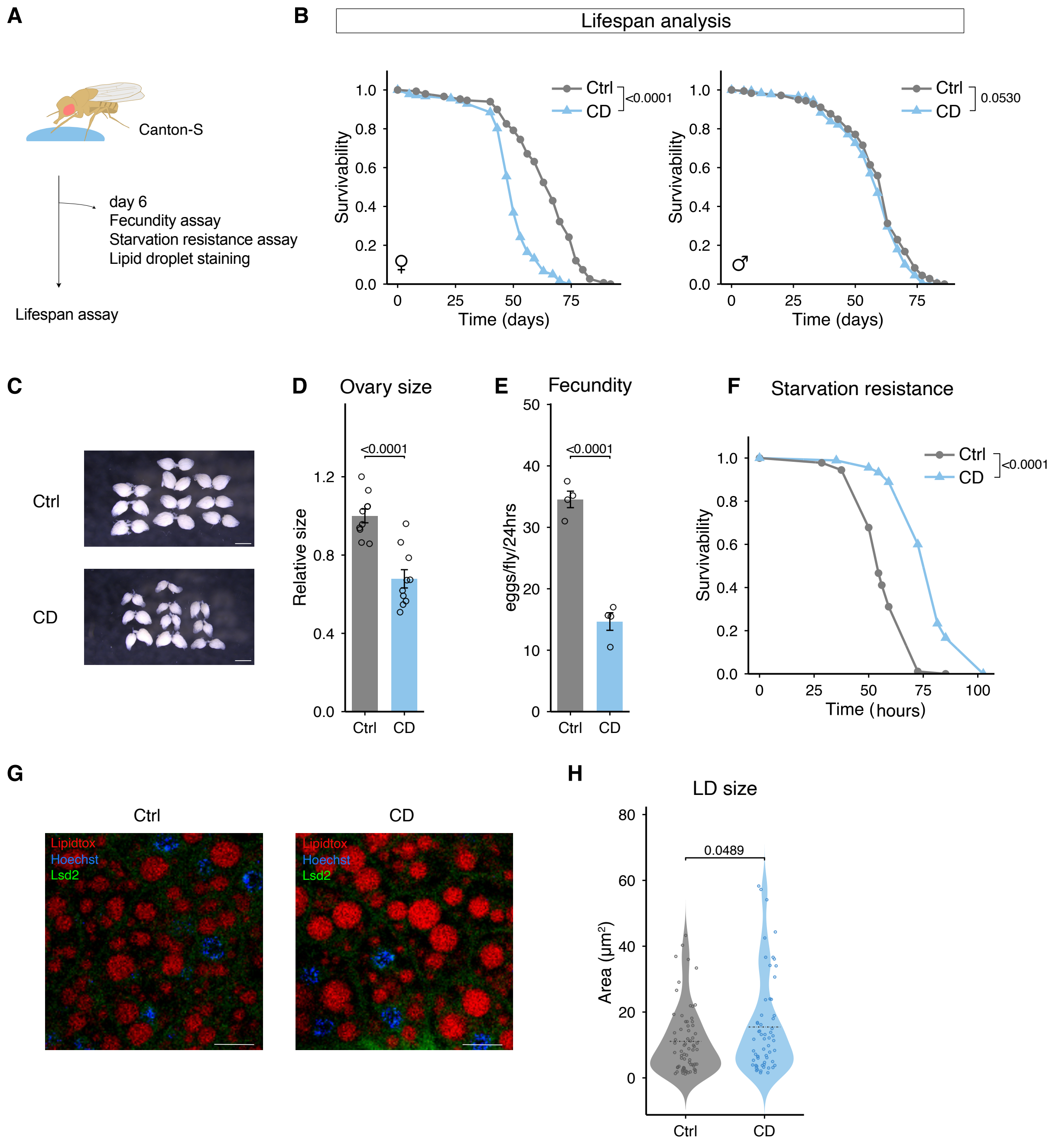


**Fig. S2. Adult choline deficient diet feeding decreases female reproductive function**

(A) Experimental scheme. Female Canton-S flies were fed with control or choline deficient diet for 6 days. (B) Lifespan analysis of male or female flies on lifelong choline deficient diet. $n$ = 149 (female Ctrl), $n$ = 179 (female CD), $n$ = 179 (male Ctrl), $n$ = 180 (male CD). (C-D) Representative image of ovaries (C) and quantification of ovary size (D) for female flies on 6 days of choline deficient diet; Scale bars, 1 mm, $n$ = 10. (E) Fecundity of female flies on 6 days of choline deficient diet; $n$ = 4. (F) Starvation resistance of female flies on 6 days of choline deficient diet; $n$ = 90. (G) Representative image of abdominal fat body of female *Lsd2-YFP* reporter flies. Scale bars, 10 μm. (H) Quantification of lipid droplet area. Data points indicate biological replicate. For (D-E, H), statistical analysis was performed by student t test. For (B,F), statistical analysis was performed by log rank test. All data are presented as mean ± s.e.m.

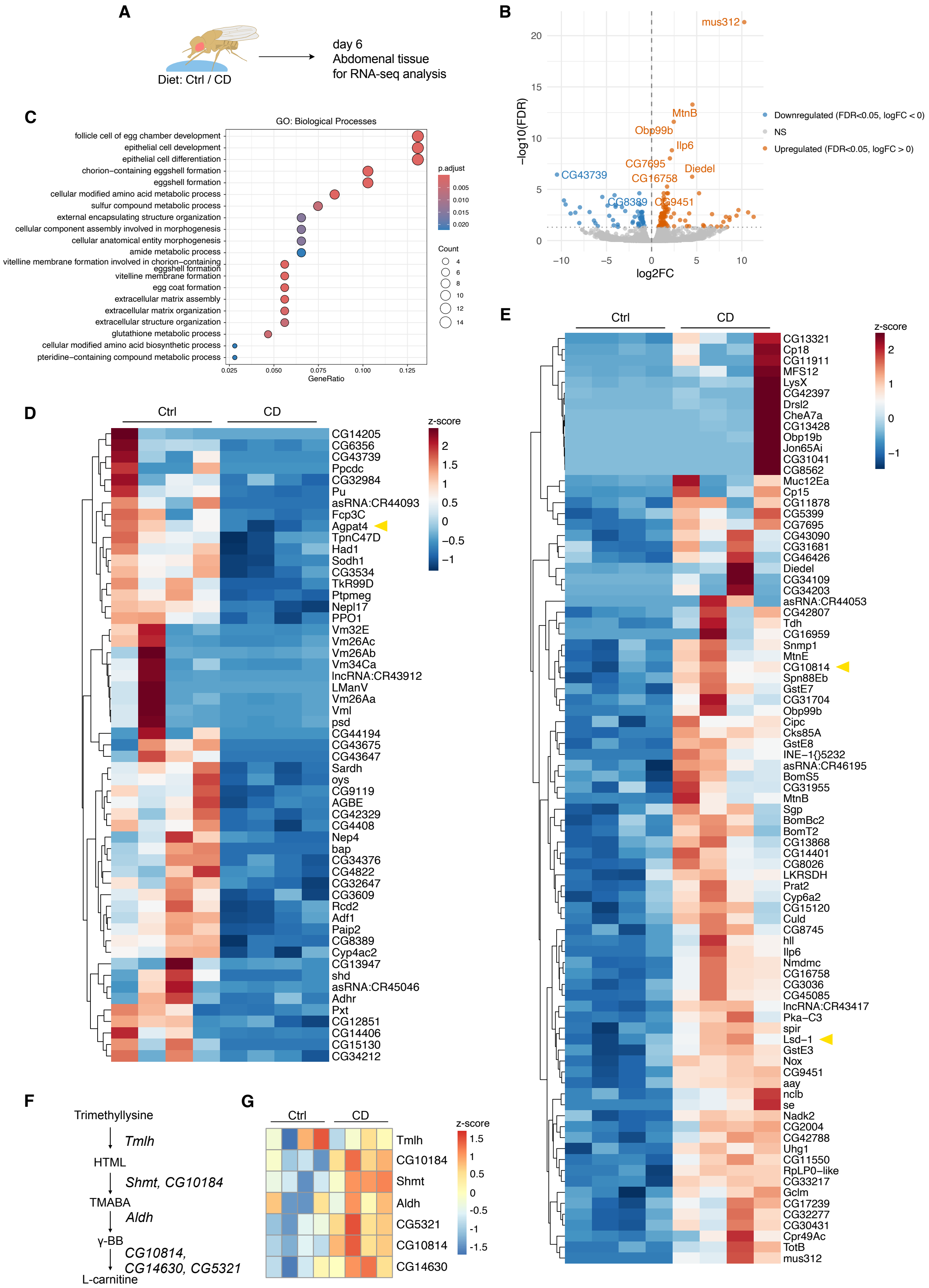
A
Diet: Ctrl / CD
day 6
Abdomenal tissue
for RNA-seq analysis
B
−log10(FDR)
log2FC
mus312
MtnB
Obp99b
Ilp6
CG7695
Diedel
CG16758
CG43739
CG8389
CG9451
Downregulated (FDR<0.05, logFC < 0)
NS
Upregulated (FDR<0.05, logFC > 0)
C
GO: Biological Processes
follicle cell of egg chamber development
epithelial cell development
epithelial cell differentiation
chorion−containing eggshell formation
eggshell formation
cellular modified amino acid metabolic process
sulfur compound metabolic process
external encapsulating structure organization
cellular component assembly involved in morphogenesis
cellular anatomical entity morphogenesis
amide metabolic process
vitelline membrane formation involved in chorion−containing eggshell formation
vitelline membrane formation
egg coat formation
extracellular matrix assembly
extracellular matrix organization
extracellular structure organization
glutathione metabolic process
cellular modified amino acid biosynthetic process
pteridine−containing compound metabolic process
GeneRatio
p.adjust
Count
D
Ctrl
CD
z-score
CG14205
CG6356
CG43739
Ppcdc
CG32984
Pu
asRNA:CR44093
Fcp3C
Agpat4
TpnC47D
Had1
Sodh1
CG3534
TkR99D
Ptpmeg
Nepl17
PPO1
Vm32E
Vm26Ac
Vm26Ab
Vm34Ca
lncRNA:CR43912
LManV
Vm26Aa
Vml
psd
CG44194
CG43675
CG43647
Sardh
oys
CG9119
AGBE
CG42329
CG4408
Nep4
bap
CG34376
CG4822
CG32647
CG3609
Rcd2
Adf1
Paip2
CG8389
Cyp4ac2
CG13947
shd
asRNA:CR45046
Adhr
Pxt
CG12851
CG14406
CG15130
CG34212
E
Ctrl
CD
z-score
CG13321
Cp18
CG11911
MFS12
LysX
CG42397
Drsl2
CheA7a
CG13428
Obp19b
Jon65Ai
CG31041
CG8562
Muc12Ea
Cp15
CG11878
CG5399
CG7695
CG43090
CG31681
CG46426
Diedel
CG34109
CG34203
asRNA:CR44053
CG42807
Tdh
CG16959
Snmp1
MtnE
CG10814
Spn88Eb
GstE7
CG31704
Obp99b
Cipc
Cks85A
GstE8
INE−1{}5232
asRNA:CR46195
BomS5
CG31955
MtnB
Sgp
BomBc2
BomT2
CG13868
CG14401
CG8026
LKRSDH
Prat2
Cyp6a2
CG15120
Culd
CG8745
hll
Ilp6
Nmdmc
CG16758
CG3036
CG45085
lncRNA:CR43417
Pka−C3
spir
Lsd−1
GstE3
Nox
CG9451
aay
nclb
se
Nadk2
CG2004
CG42788
Uhg1
CG11550
RpLP0−like
CG33217
Gclm
CG17239
CG32277
CG30431
Cpr49Ac
TotB
mus312
F
Trimethyllysine
Tmlh
HTML
Shmt, CG10184
TMABA
Aldh
γ-BB
CG10814, CG14630, CG5321
L-carnitine
G
Ctrl
CD
z-score
Tmlh
CG10184
Shmt
Aldh
CG5321
CG10814
CG14630

**Fig. S3. Transcriptome analysis of adult choline deficiency**

(**A**) Experimental scheme. RNA-seq analysis was performed on abdominal tissue of female Canton-S flies fed with control or choline deficient diet for 6 days. (**B**) Volcano plot. Upregulated genes (FDR<0.05, logFC>0) is shown in orange, and downregulated genes (FDR<0.05, logFC<0) is shown in blue. (**C**) Gene ontology analysis for biological processes of DEG (FDR<0.05). (**D**) Heatmap showing z scores for DEGs that were downregulated by choline deficient diet (FDR<0.05, logFC<0). Yellow arrow shows genes related to lipid metabolism. (**E**) Heatmap showing z scores for DEGs that were upregulated by choline deficient diet (FDR<0.05, logFC>0). Yellow arrow shows genes especially related to lipid metabolism. (**F**) Schematics of predicted carnitine biosynthesis pathway. (**G**) Heatmap showing z scores of genes involved in carnitine biosynthesis pathway.

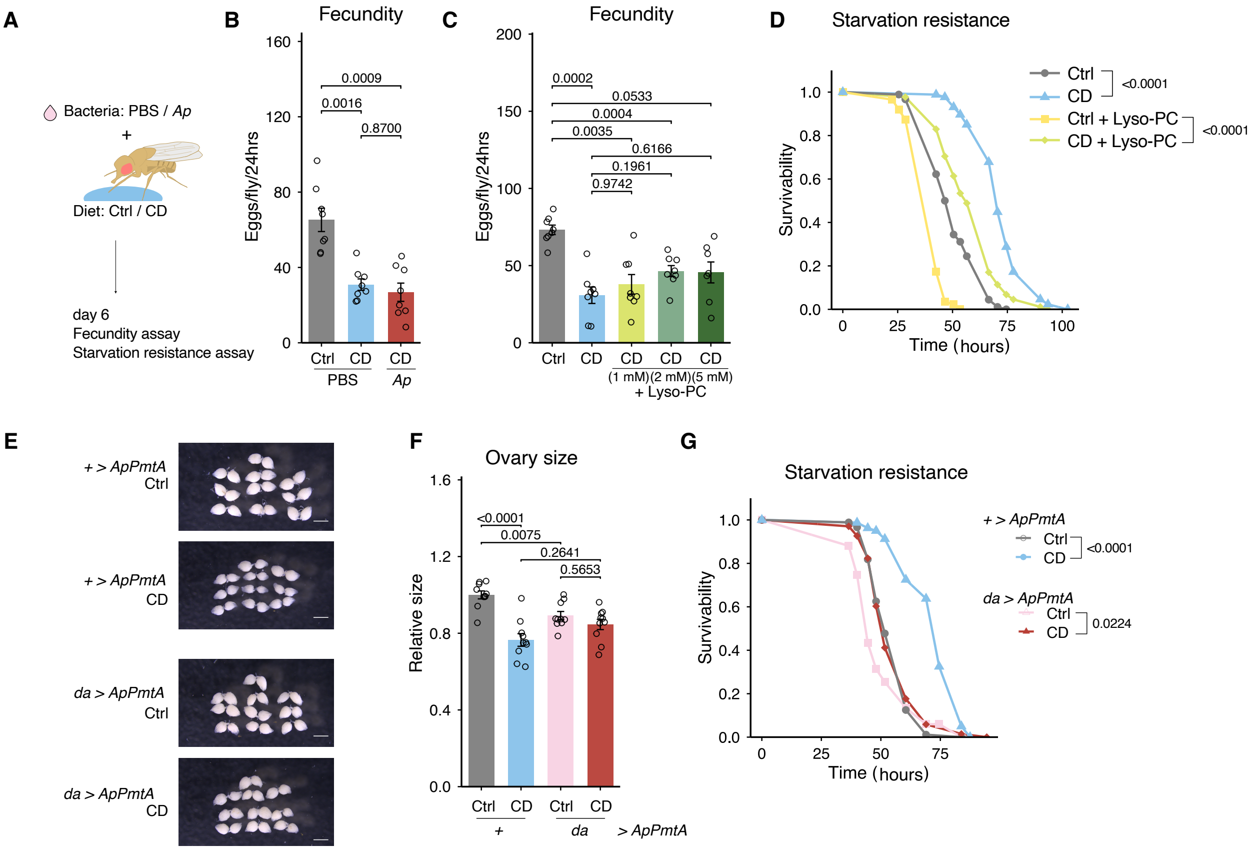

**Fig. S4. Effects of adult choline deficiency cannot be rescued by Ap or Lyso-PC supplementation, but just by *PmtA* overexpression** (**A**) Experimental scheme. *Ap* was cultured separately and supplemented to choline deficient diet and given to the flies. (**B**) Fecundity after 6 days on *Ap* supplemented choline deficient diet; $n = 8$. (**C**) Fecundity after 6 days on choline deficient diet supplemented with Lyso-PC; $n = 8$. (**D**) Starvation resistance after 6 days on choline deficient diet supplemented with 0.5 mM Lyso-PC; $n = 90$ (Ctrl), $n = 87$ (CD), $n = 87$ (Ctrl+Lyso-PC), $n = 88$ (CD+Lyso-PC). (**E-F**) Representative image of ovaries (**E**) and quantification of ovary size (**F**); $n = 10$. (**G**) Starvation resistance for *da>ApPmtA* flies after 6 days on choline deficient diet; $n = 88$ (Ctrl_+>*ApPmtA*), $n = 80$ (CD_+>*ApPmtA*), $n = 67$ (Ctrl_*da>ApPmtA*), $n = 68$ (CD_*da>ApPmtA*). Data points indicate biological replicate. For (**B**, **C** and **F**), statistical analysis were performed by one way analysis of variance (ANOVA). The *p* values were determined by post hoc analysis with Šídák's multiple comparison test. For (D,G), statistical analysis was performed by log rank test. All data are presented as mean ± s.e.m.

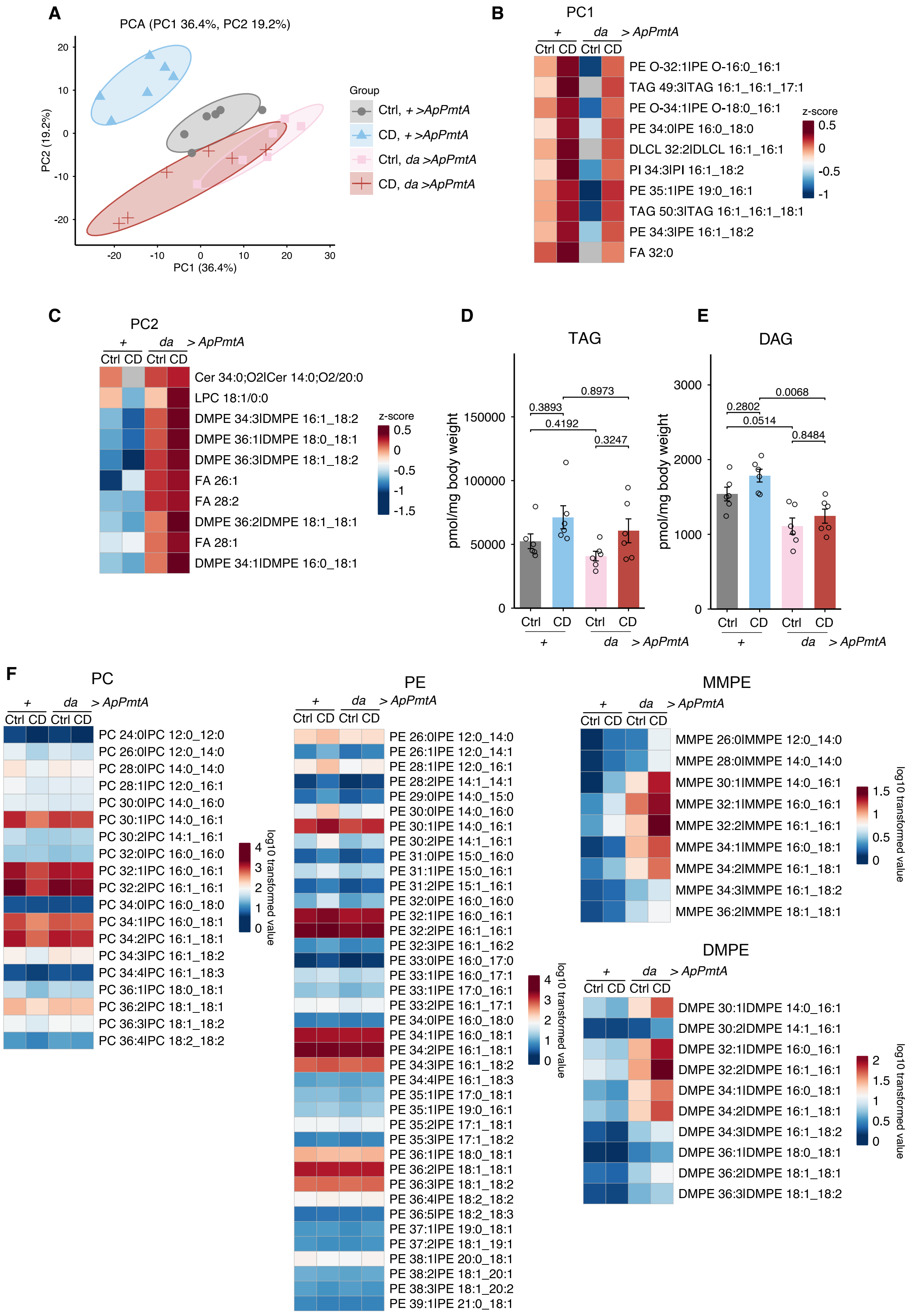
A
PCA (PC1 36.4%, PC2 19.2%)
PC2 (19.2%)
PC1 (36.4%)
Group
Ctrl, + >ApPmtA
CD, + >ApPmtA
Ctrl, da >ApPmtA
CD, da >ApPmtA
B
PC1
+ da > ApPmtA
Ctrl CD Ctrl CD
PE O-32:1|PE O-16:0_16:1
TAG 49:3|TAG 16:1_16:1_17:1
PE O-34:1|PE O-18:0_16:1
PE 34:0|PE 16:0_18:0
DLCL 32:2|DLCL 16:1_16:1
PI 34:3|PI 16:1_18:2
PE 35:1|PE 19:0_16:1
TAG 50:3|TAG 16:1_16:1_18:1
PE 34:3|PE 16:1_18:2
FA 32:0
z-score
C
PC2
Cer 34:0;O2|Cer 14:0;O2/20:0
LPC 18:1/0:0
DMPE 34:3|DMPE 16:1_18:2
DMPE 36:1|DMPE 18:0_18:1
DMPE 36:3|DMPE 18:1_18:2
FA 26:1
FA 28:2
DMPE 36:2|DMPE 18:1_18:1
FA 28:1
DMPE 34:1|DMPE 16:0_18:1
D
TAG
pmol/mg body weight
0.3893
0.8973
0.4192
0.3247
E
DAG
0.2802
0.0068
0.0514
0.8484
F
PC
PC 24:0|PC 12:0_12:0
PC 26:0|PC 12:0_14:0
PC 28:0|PC 14:0_14:0
PC 28:1|PC 12:0_16:1
PC 30:0|PC 14:0_16:0
PC 30:1|PC 14:0_16:1
PC 30:2|PC 14:1_16:1
PC 32:0|PC 16:0_16:0
PC 32:1|PC 16:0_16:1
PC 32:2|PC 16:1_16:1
PC 34:0|PC 16:0_18:0
PC 34:1|PC 16:0_18:1
PC 34:2|PC 16:1_18:1
PC 34:3|PC 16:1_18:2
PC 34:4|PC 16:1_18:3
PC 36:1|PC 18:0_18:1
PC 36:2|PC 18:1_18:1
PC 36:3|PC 18:1_18:2
PC 36:4|PC 18:2_18:2
log10 transformed value
PE
PE 26:0|PE 12:0_14:0
PE 26:1|PE 12:0_14:1
PE 28:1|PE 12:0_16:1
PE 28:2|PE 14:1_14:1
PE 29:0|PE 14:0_15:0
PE 30:0|PE 14:0_16:0
PE 30:1|PE 14:0_16:1
PE 30:2|PE 14:1_16:1
PE 31:0|PE 15:0_16:0
PE 31:1|PE 15:0_16:1
PE 31:2|PE 15:1_16:1
PE 32:0|PE 16:0_16:0
PE 32:1|PE 16:0_16:1
PE 32:2|PE 16:1_16:1
PE 32:3|PE 16:1_16:2
PE 33:0|PE 16:0_17:0
PE 33:1|PE 16:0_17:1
PE 33:1|PE 17:0_16:1
PE 33:2|PE 16:1_17:1
PE 34:0|PE 16:0_18:0
PE 34:1|PE 16:0_18:1
PE 34:2|PE 16:1_18:1
PE 34:3|PE 16:1_18:2
PE 34:4|PE 16:1_18:3
PE 35:1|PE 17:0_18:1
PE 35:1|PE 19:0_16:1
PE 35:2|PE 17:1_18:1
PE 35:3|PE 17:1_18:2
PE 36:1|PE 18:0_18:1
PE 36:2|PE 18:1_18:1
PE 36:3|PE 18:1_18:2
PE 36:4|PE 18:2_18:2
PE 36:5|PE 18:2_18:3
PE 37:1|PE 19:0_18:1
PE 37:2|PE 18:1_19:1
PE 38:1|PE 20:0_18:1
PE 38:2|PE 18:1_20:1
PE 38:3|PE 18:1_20:2
PE 39:1|PE 21:0_18:1
MMPE
MMPE 26:0|MMPE 12:0_14:0
MMPE 28:0|MMPE 14:0_14:0
MMPE 30:1|MMPE 14:0_16:1
MMPE 32:1|MMPE 16:0_16:1
MMPE 32:2|MMPE 16:1_16:1
MMPE 34:1|MMPE 16:0_18:1
MMPE 34:2|MMPE 16:1_18:1
MMPE 34:3|MMPE 16:1_18:2
MMPE 36:2|MMPE 18:1_18:1
DMPE
DMPE 30:1|DMPE 14:0_16:1
DMPE 30:2|DMPE 14:1_16:1
DMPE 32:1|DMPE 16:0_16:1
DMPE 32:2|DMPE 16:1_16:1
DMPE 34:1|DMPE 16:0_18:1
DMPE 34:2|DMPE 16:1_18:1
DMPE 34:3|DMPE 16:1_18:2
DMPE 36:1|DMPE 18:0_18:1
DMPE 36:2|DMPE 18:1_18:1
DMPE 36:3|DMPE 18:1_18:2

**Fig. S5.** ***ApPmtA*** **overexpression rescues the lipidomic profile of choline deficiency**
(**A**) Principal Component Analysis of whole body lipidome analysis for *da>ApPmtA* flies on choline deficient diet. (**B-C**) Top 10 genes that contributed to Principal Component 1 (**B**) and 2 (**C**). (**D-E**) Quantification of total TAG (**D**) and DAG (**E**) levels. (**F**) Heatmap showing log 10 transformed concentration for metabolites on the methylation pathway. For (D and E), statistical analysis was performed by one way analysis of variance (ANOVA). The *p* values were determined by post hoc analysis with Šídák's multiple comparison test. All data are presented as mean ± s.e.m.

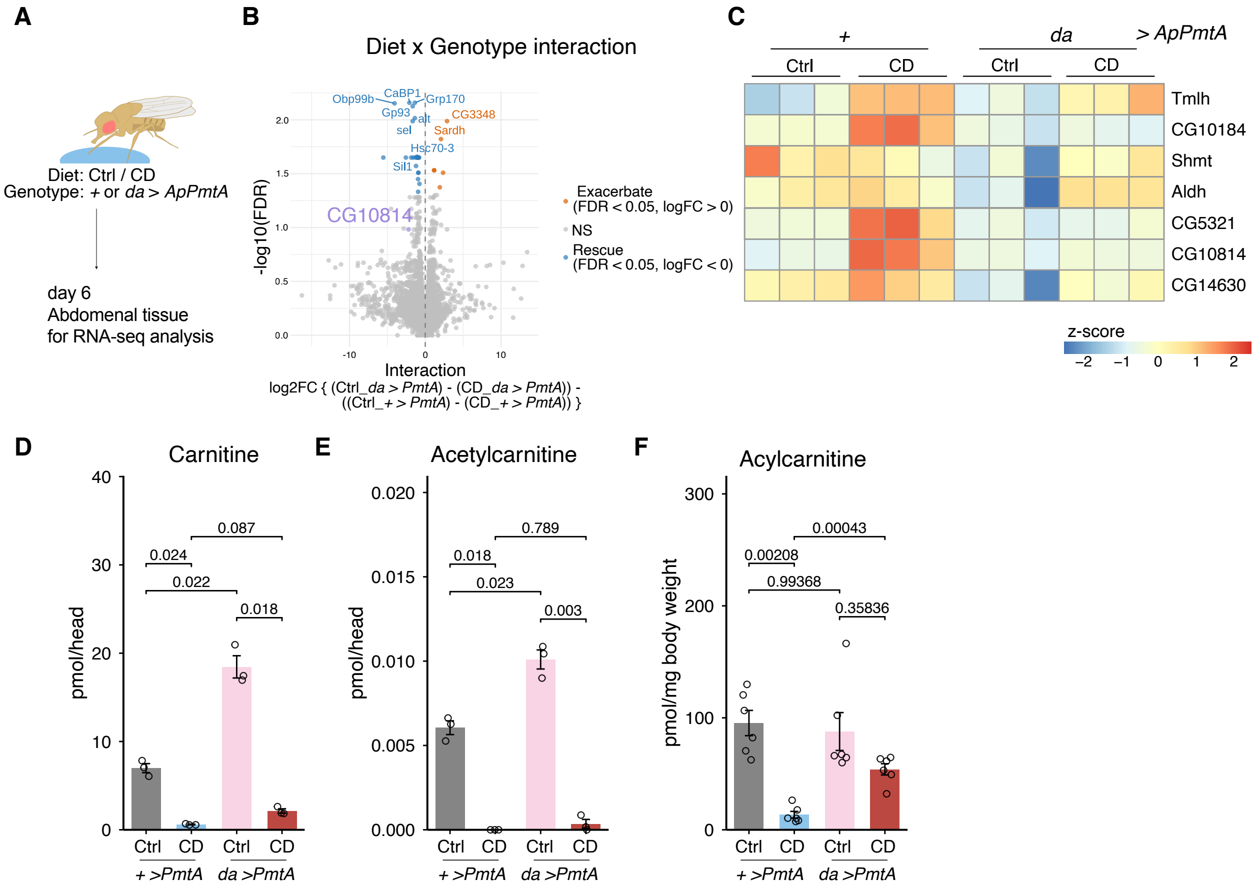


**Fig. S6.** ***ApPmtA c*****an partially rescue the altered carnitine metabolism induced by choline deficiency**

(**A**) Experimental scheme. RNA-seq analysis was performed on abdominal tissue of female *da>ApPmtA* flies fed with control or choline deficient diet for 6 days. (**B**) Volcano plot. Upregulated genes (FDR<0.05, logFC>0) are shown in orange, downregulated genes (FDR<0.05, logFC<0) are shown in blue, and *CG10814* is shown in purple. (**C**) Heatmap showing z scores of genes involved in carnitine biosynthesis pathway. (**D-E**) Quantification of carnitine (D) and acetyl carnitine (E) in head samples of *da>ApPmtA* flies fed with control or choline deficient diet for 6 days. (**F**) Quantification of long-chain acylcarnitine in whole body samples of *da>ApPmtA* flies fed with control or choline deficient diet for 6 days. For (D-F), statistical analysis were performed by one way analysis of variance (ANOVA). The *p* values were determined by post hoc analysis with Šídák's multiple comparison test. All data are presented as mean ± s.e.m.

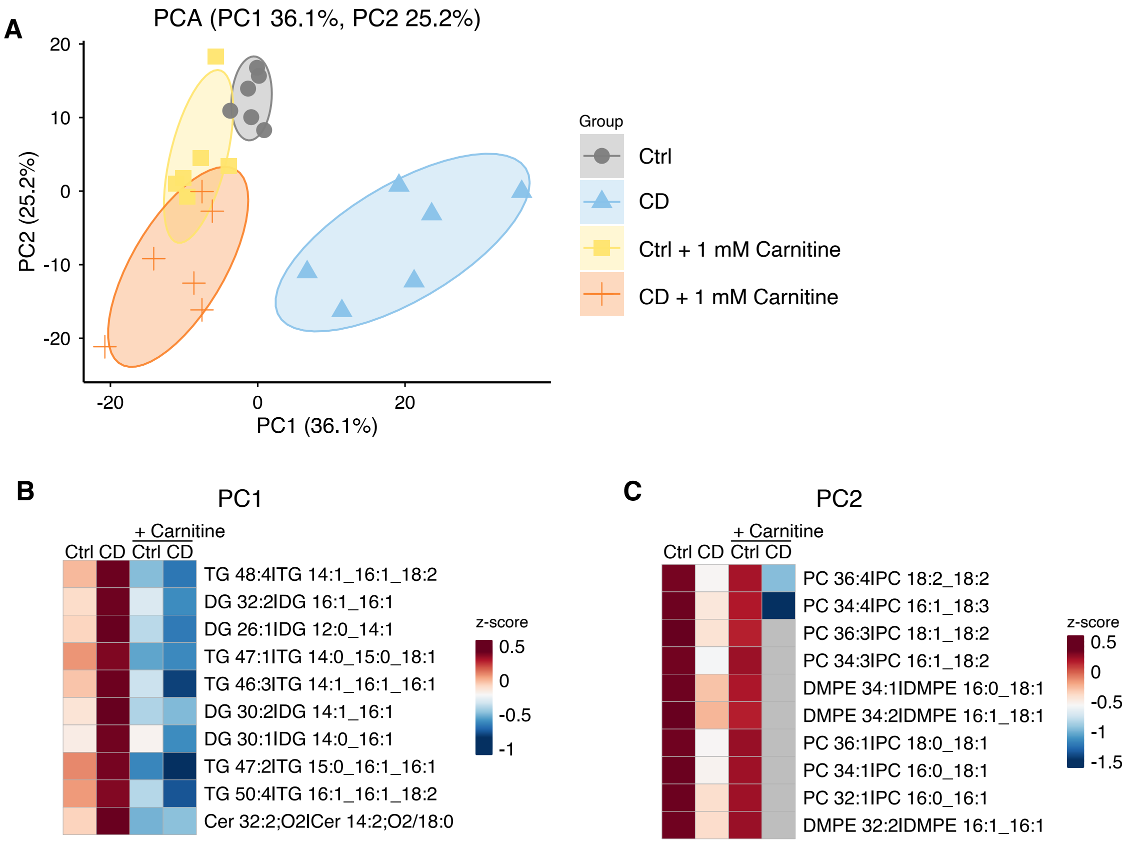


**Fig. S7. Lipidome analysis for whole body of flies on CD supplemented with carnitine**

(**A**) Principal Component Analysis of whole body lipidome analysis for flies on choline deficient diet supplemented with carnitine. (**B-C**) Heatmap of z score for top 10 lipids that contributed to Principal Component 1 (**B**) and 2 (**C**).

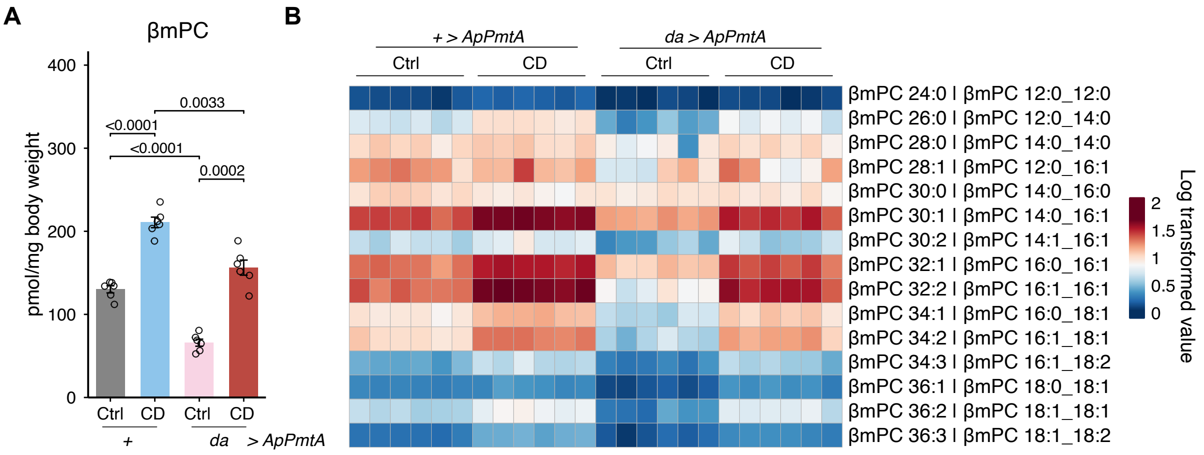


**Fig. S8. βmPC is increased under CD in *PmtA* overexpressing flies**

(A) Quantification of βmPC in the whole body of *da>ApPmtA* flies on CD. (B) All species of annotated βmPC. Heatmap shows log 10 transformed values of βmPC. For (A), statistical analysis were performed by one way analysis of variance (ANOVA). The p values determined by post hoc analysis with Šídák's multiple comparison test. All data are presented as mean ± s.e.m.

| | ln(HR) | HR | se(HR) | P value (zscore) |
|---|---|---|---|---|
| **CD (diet)** | 1.4303 | 4.1798 | 0.1727 | <0.0001*** |
| **Lyso-PC (diet)** | -1.708 | 0.1812 | 0.1709 | <0.0001*** |
| **CD (diet)* Lyso-PC (diet)** | -0.5161 | 0.5969 | 0.2286 | 0.024 |

**Table S1. Effects of dietary choline and lyso-PC on hazard ratios of mortality**

| | ln(HR) | HR | se(HR) | P value (zscore) |
|---|---|---|---|---|
| **CD (diet)** | 2.7747 | 16.0336 | 0.1721 | <0.0001*** |
| ***ApPmtA* overexpression** | -0.5340 | 0.5863 | 0.1231 | <0.0001*** |
| **CD (diet)* *ApPmtA*** | -2.8548 | 0.0576 | 0.2136 | <0.0001*** |

**Table S2. Effects of dietary choline and *ApPmtA* overexpression on hazard ratios of mortality**

| | ln(HR) | HR | se(HR) | P value (zscore) |
|---|---|---|---|---|
| **CD (diet)** | -1.2522 | 0.2859 | 0.1687 | <0.0001*** |
| ***ApPmtA* overexpression** | 0.2026 | 1.2246 | 0.1657 | 0.221314 |
| **CD (diet)* *ApPmtA*** | 0.8234 | 2.2782 | 0.2430 | <0.0001*** |

**Table S3. Effects of dietary choline and *ApPmtA* overexpression on hazard ratios of mortality under starvation stress**

| | ln(HR) | HR | se(HR) | P value (zscore) |
|---|---|---|---|---|
| **CD (diet)** | 0.7726 | 2.1653 | 0.1142 | <0.0001*** |
| **Carnitine supplementation (diet)** | 0.6808 | 1.9755 | 0.1083 | <0.0001*** |
| **CD (diet)* Carnitine supplementation (diet)** | -0.9742 | 0.3775 | 0.1574 | <0.0001*** |

**Table S4. Effects of dietary choline and carnitine on hazard ratios of mortality**

| | ln(HR) | HR | se(HR) | P value (zscore) |
|---|---|---|---|---|
| **CD (diet)** | 0.7897 | 2.2028 | 0.1120 | <0.0001*** |
| **βmc supplementation (diet)** | 0.0788 | 1.0820 | 0.1122 | <0.0001*** |
| **CD (diet)*βmc supplementation (diet)** | -0.6063 | 0.5454 | 0.1561 | <0.0001*** |

**Table S5. Effects of dietary choline and β-methylcholine on hazard ratios of mortality**

**Data S1. (separate file)**
LSI checklist for bacterial samples.

**Data S2. (separate file)**
LSI checklist for fly samples.
